# FOXA1 preserves cell polarity and restrains lysosome biogenesis in non-small cell lung adenocarcinoma

**DOI:** 10.64898/2026.04.09.717383

**Authors:** Xuecong Wang, Bingwen Zhang, Chu Sun, Meiyu Huang, Weitao Huang, Binghao Zhang, Xuan Zhang, Xia Ren, Ling Luo, Hengrui Liang, Youtao Zhou, Guifa Zhong, Shouheng Lin, Micky D. Tortorella, Tuan Zea Tan, Wenhua Liang, Jean Paul Thiery, Jianxing He

## Abstract

**Background:** This study investigates the role of the pioneer transcription factor FOXA1 as a master gene in sustaining epithelial cell polarization in early-stage lung adenocarcinoma. The partial loss of FOXA1 is explored to determine if it will affect plasticity and progression of lung adenocarcinoma. The study also addresses the transcriptional circuitry that links polarity defects to lysosome homeostasis.

**Methods:** A multiomics approach was used to define the status of the chromatin in epithelial and mesenchymal states of A549 adenocarcinoma cells obtained with a newly synthetized TGF-β receptor inhibitor or TGF-β respectively.

The study leveraged ATAC-seq, RNA sequencing, Cut&Tag sequencing of FOXA1 and histone marks profiling. The functional impact of FOXA1 was examined by partial silencing in vitro and by heterozygous FOXA1 deletion in a *Kras*^G12D^ mouse model. Three-dimensional organoid culture, high-resolution electron microscopy, spatial transcriptomics and multiplex immunohistochemistry assessed carcinoma cell polarity, proliferation, the tumor microenvironment and organelle content. Group differences were evaluated with two-tailed t tests or one-way analysis of variance.

**Results:** FOXA1 binding and expression were highest in cells harboring an epithelial phenotype.

In mouse *Kras*^G12D^ LUAD tumors FOXA1 marked polarized, CDH1-positive cells; heterozygous loss diminished CDH1, disrupted apical-basal architecture, lowered organoid-forming efficiency and remodeled the immune microenvironment. Spatial transcriptomics and ultrastructural analyses showed that FOXA1-deficient carcinoma cells accumulated lysosomes, down-regulated vesicle fusion genes of the SNARE family and activated the lysosomal CLEAR gene network. FOXA1 occupied enhancers of lysosome-associated genes and competed with the transcription factor TFE3, thereby suppressing transcription of cathepsin B and cathepsin C and restricting lysosome biogenesis.

**Conclusions:** FOXA1 is a central regulator that preserves epithelial cell polarity and limits lysosome formation in lung adenocarcinoma. Targeting the FOXA1–TFE3–lysosome axis may affect tumor plasticity and provide new therapeutic opportunities.

## Background

Lung adenocarcinoma, the most prevalent subtype of non-small cell lung cancer (NSCLC), is a highly heterogeneous disease characterized by cell plasticity and complex tumor microenvironment (TME) interactions[1, 2]. Epithelial-mesenchymal transition (EMT) is a pivotal process that drives tumor progression, metastasis, and therapeutic resistance, with the dynamic regulation of cell polarity and phenotype underpinning these malignant features[3]. Transforming growth factor-β (TGF-β) is a predominant EMT driver in lung adenocarcinoma, and pharmacological or genetic blockade of the pathway can re-establish an epithelial phenotype[4, 5]. However, the molecular circuitry that stabilizes epithelial identity and polarity once TGF-β signaling is curtailed remains incompletely understood.

Recent studies have highlighted the importance of lineage-specific transcription factors in maintaining epithelial characteristics and suppressing EMT in lung cancer[6, 7]. Among these, the Forkhead box protein A1 (FOXA1) has emerged as a central regulator of epithelial differentiation, playing critical roles in lung development, tissue homeostasis, and cancer progression[8–10]. Furthermore, FOXA1 has also been proven to be the key regulator working with FOXA2 to determine cancer cell lineage in lung adenocarcinoma[11, 12], yet its function in restraining lung cancer cell plasticity is still unresolved.

Concurrently, lysosomes, classically known for their role in cellular degradation and recycling, have emerged as crucial hubs for regulating metabolism, cell plasticity and invasion in cancer[13, 14]. Their biogenesis is controlled by the Coordinated Lysosomal Expression and Regulation (CLEAR) gene network, which is orchestrated by the MiT/TFE family of transcription factors, such as TFE3[15, 16]. Although the role of lysosomes in cancer is a subject of intense research, a direct link between their function and cell identity regulators like FOXA1 has not been previously well established[17, 18].

In this study, we investigate the role of FOXA1 in maintaining epithelial cell polarity and suppressing lysosome biogenesis in lung adenocarcinoma. A combination of in vitro cell line models, genetically engineered mouse models, and advanced omics approaches, including ATAC-seq, RNA-seq, CUT&Tag sequencing, and spatial transcriptomics, enabled us to comprehensively characterize its functions. Our findings reveal that FOXA1 is highly enriched in polarized, well-differentiated carcinoma cells and is essential for sustaining epithelial cell identity, robust proliferation, and tumorigenic potential. Mechanistically, FOXA1 antagonizes TGF-β-mediated EMT, restricts lysosome biogenesis through competitive inhibition of TFE3 at lysosomal gene loci, leading to reduced protein production in polarized lung adenocarcinoma region.

Collectively, our work uncovers a previously unrecognized link between FOXA1, cell polarity, and lysosome homeostasis in lung adenocarcinoma, providing new insights into the molecular regulation of tumor progression and offering potential avenues for therapeutic intervention.

## Methods

### Kinome Selectivity

Kinase selectivity was assessed at Eurofins Pharma Discovery Services (Dundee, UK) on the KinaseProfiler® radiometric platform following the vendor’s SOPs. B1.14 was dissolved in anhydrous DMSO to 10 mM and diluted 50-fold in assay buffer to create a 50 X working stock. Appropriate volumes of this stock were transferred to 96-well plates, maintaining a final DMSO concentration of ≤ 2% (v/v). A premixed solution of recombinant kinase and its cognate peptide substrate was then added, and reactions were initiated by the addition of ATP at the concentration specified by Eurofins; the compound was not pre-incubated with the enzyme/substrate mixture. Reactions proceeded for the time recommended in the KinaseProfiler protocol and were terminated according to standard procedures. Kinase activity in the presence of B1.14 was calculated as [(C_sample – C_no-enzyme)/(C_plus-enzyme – C_no-enzyme)] × 100, where C denotes scintillation counts. All measurements were performed in duplicate unless otherwise stated.

### Tissue Collection and Paraffin Embedding

Mouse lung tissue samples were collected in accordance with the ethical approval procedures (IACUC No. S2022-508). The tissues were immediately fixed in 4% paraformaldehyde (PFA) for 24 hours at room temperature. Following fixation, the samples were dehydrated and cleared using an automated tissue processor (HistoCore Pearl, Leica). The tissues were then embedded in paraffin blocks using a paraffin embedding station (HistoCore Arcadia H+C, Leica) and allowed to solidify at room temperature. The paraffin blocks were stored at 4°C until they were sectioned.

For histological analysis, paraffin-embedded tissues were sectioned at a thickness of 4 µm using a rotary microtome (HistoCore Autocut, Leica). Slides were dried to ensure tissue adherence before staining.

### IHC staining

Endogenous peroxidase activity was blocked by incubating the sections in 3% hydrogen peroxide for 10 minutes at room temperature. Non-specific binding was minimized by incubating the sections with 5% bovine serum albumin (BSA) in phosphate-buffered saline (PBS) for 30 minutes. Primary antibodies specific to the target protein were applied and incubated overnight at 4°C. After washing with PBS, the sections were incubated with a horseradish peroxidase (HRP)-conjugated secondary antibody (dilution 1:200) for 1 hour at room temperature.

The antigen-antibody complexes were visualized using 3,3’-diaminobenzidine (DAB) as the chromogen, and the reaction was monitored under a microscope. The sections were counterstained with hematoxylin for 1 minute, followed by dehydration through a graded series of ethanol and xylene. Finally, the slides were mounted with a coverslip using a permanent mounting medium.

Images of stained sections were captured using a digital pathology scanner (Aperio CS2, Leica). Quantitative analysis of staining intensity was performed using HALO software (Indica Labs, USA).

### MIHC (TSA) staining and analysis

To identify cell subsets within human or mouse paraffin sections, a multiplex immunohistochemistry (mIHC) staining protocol was performed using the PANO 7-plex IHC kit (Cat. No. 0004100100, Panovue). Sequential application of different primary antibodies was followed by incubation with horseradish peroxidase (HRP)-conjugated secondary antibodies and subsequent tyramide signal amplification (TSA). Microwave treatment was employed after each TSA step to enhance antigen retrieval. Once all target antigens were labeled, nuclei were counterstained with 4′,6-diamidino-2-phenylindole (DAPI, Sigma-Aldrich). Whole-slide fluorescent imaging was performed using an Olympus VS200 scanner (Olympus) equipped with an Olympus UPLXAPO 20x objective lens. The acquired fluorescence images were analyzed using the HALO software (Indica Labs).

### Bulk RNA-seq Sample Preparation

Approximately 5 × 10^5^ cells were harvested, and total RNA was isolated using TRIzol reagent. Genomic DNA was digested with DNase I. The integrity and purity of RNA were evaluated using an Agilent Bioanalyzer 2100. Poly(A)+ mRNA was enriched using oligo(dT) magnetic beads, and sequencing libraries were constructed with the Illumina TruSeq Stranded mRNA Library Prep Kit, following the manufacturer’s protocols. Library quality was verified by Qubit quantification and Bioanalyzer analysis. Final libraries were sequenced on an Illumina NovaSeq 6000 platform, generating 150 bp paired-end reads.

### Bulk mRNA-seq Data Processing and Analysis

The paired-end RNA-Seq reads were trimmed of adapters, and low-quality reads were removed by Cutadapt(v2.8)[19] with the option of ‘-m 20 -q 20’. Trimmed data were aligned to the GRCh38/hg38 (for human) or GRCm38/mm10 (for mouse) reference genomes using HISAT2 (v2.1.0)[20] with the settings of ‘-p 20 -t --mm --no-unal--no-mixed --no-discordant’. Gene expression levels were quantified for each sample in terms of TPM (Transcripts Per Million) and FPKM (Fragments Per Kilobase of transcript per Million mapped reads) using the rnanorm Python package (v2.1.0) via the functions TPM() and FPKM(). For differential expression analysis, raw read counts were obtained using featureCounts (v2.0.0)[21]. Differential expression was analyzed with the R/Bioconductor package DESeq2 (v1.26.0)[22]. Genes with a false discovery rate (FDR) below 0.05 and an absolute fold change greater than 1.5 were considered differentially expressed (DE). Heatmaps were generated using z-score transformed TPM values with the Python package PyComplexHeatmap (v1.6)[23]. Functional enrichment analyses, including Gene Ontology (GO) and KEGG pathway annotations, were performed with the Python package GSEApy (v1.0.6)[24].

### ATAC-seq and Cut&Tag Library Preparation

ATAC-seq library preparation was conducted using the TruePrep DNA Library Prep Kit V2 for Illumina (TD501, Vazyme), employing a transposase-based library construction method following the manufacturer’s instructions. In brief, 5 × 10^4^ cell samples were collected. Cell lysis was performed to isolate intact nuclei, which were subsequently counted. During this stage, organelle DNA was efficiently removed. Tn5 transposase was then introduced to fragment the DNA in its open state. The resulting DNA fragments were recovered, ligated with adapters to construct a complete library, and subjected to PCR amplification using TruePrep Index Kit V2 for Illumina (TD202, Vazyme). Following amplification, the library underwent subsequent purification using VAHTS DNA Clean Beads (Vazyme #N411) and ready for quality control and sequencing.

For the preparation of Cut&Tag-seq libraries, the Hyperactive Universal CUT&Tag Assay Kit for Illumina (TD903, Vazyme) was utilized in accordance with the manufacturer’s protocol. In brief, 1.0 × 10^5^ cells were gently resuspended in 10 μL of binding buffer and subsequently conjugated to pre-activated concanavalin A-coated magnetic beads. The bead-cell complexes were incubated overnight at 4°C in 50 μL of antibody buffer containing primary antibody, followed by incubation with a secondary antibody the next day. DNA tagmentation was performed using the pA/G-Tn5 adapter complex. The tagmented DNA was then isolated using DNA extraction beads and amplified through PCR using the TruePrep Index Kit V2 for Illumina (TD202, Vazyme). Finally, the library was purified with VAHTS DNA Clean Beads (Vazyme #N411) and prepared for quality assessment and sequencing.

### ATAC-seq and CUT&Tag Data Processing and Analysis

For ATAC-seq and CUT&Tag data, adapter sequences were trimmed, and low-quality reads were filtered using Cutadapt (v2.8). The trimmed reads were aligned to the GRCh38/hg38 (for human) or GRCm38/mm10 (for mouse) using Bowtie2 (2.3.5.1)[25] with the settings ‘–mm, --very-sensitive, --no-unal, --no-mixed, --no-discordant’. Sambamba (v0.7.1)[26] was used to mark low-quality mapped reads (with a mapping quality threshold of ‘-mapq 3’). Duplicate reads were identified and marked using markdup, and subsequently, both low-quality and duplicate reads were removed. Reads mapped to mitochondrial DNA (chrM) were excluded using grep -v ‘chrM’. Peaks were called using MACS2 (v2.2.6)[27] with the settings ‘-q 0.001, -f BAMPE’, ‘-g hs’ for human or ‘-g mm’ for mouse, and the ‘--call-summits’ option enabled. Peaks were annotated using HOMER (v4.9.1)[28] annotatePeaks.pl tools. De novo Motif discovery was performed using HOMER (v4.9.1) findMotifsGenome.pl tools with the default option. BigWig files were generated using deepTools (v3.5.1)[29] bamCoverage with the parameters ‘--ignoreDuplicates’ and ‘--normalizeUsing PRGC’. Peak density pileups were calculated using deepTools computeMatrix (reference-point mode) with ‘--skipZeros’ and ‘--missingDataAsZero’ options. Visualization of data was performed using deepTools plotHeatmap with ‘--sortUsing sum’, and plotProfile with the ‘--perGroup’ option. Gene ontology (GO), KEGG pathway and gene expression measures were called by first collecting all TSSs within 50 kb of an ATAC-seq or CUT&Tag peak and then performing enrichment analysis with GSEApy (v1.0.6) or measuring gene expression. BEDTools[30] and Pybedtools[31] were used to determine genomic peak intersections, proximity analysis, and other similar calculations.

### Differential Analysis of ATAC-seq and CUT&Tag Data

Differential accessibility (for ATAC-seq) or differential binding (for CUT&Tag) between libraries of each data type was assessed using the CSAW (ChIP-Seq Analysis with Windows) package (v1.32.0)[32]. Biological replicates were maintained separately for the analysis to account for variability between samples. For each data type, read parameters were defined using the readParam function with the options ‘pe=both’ (for paired-end reads) and ‘Max.frag=1000’ (for maximum fragment length). Reads mapping to blacklisted genomic regions were excluded from the analysis. Average log counts per million (LogCPM) were computed using the ‘aveLogCPM()’ function, and low-abundance windows were filtered out based on a predefined threshold across different libraries. For transcription factors and the histone marker H3K27ac, sliding windows with a width of 600 bp were used, a 2 kb neighborhood around each window was defined to estimate local background levels. Read counts within these neighborhoods were stored using the ‘wider()’ function. The ‘filterWindowsLocal()’ function was then applied to filter out non-enriched features, retaining only those windows that exhibited significant local enrichment with at least a 4-fold change. To aggregate locally enriched windows, larger sliding windows of 10,000 bp were used. TMM normalization was performed using the ‘normFactors()’ function to adjust for library size differences. Differential binding analysis was conducted using the quasi-likelihood (QL) framework in the edgeR package (v3.40.2)[33], with the ‘glmQLFit()’ function set to ‘robust=TRUE’ to ensure model robustness. The design matrix was constructed to incorporate experimental conditions, including FOXA1 knockdown or drug treatments. Differentially tested windows were merged into regions using the ‘mergeWindows()’ function, with a tolerance distance between windows set to ‘tol=500L’ and a maximum merged window size of ‘max.width=5000L’. To identify the most significant window within each merged region, the ‘getBestTest()’ function was used, providing a representative p-value and false discovery rate (FDR) for each merged window.

### TOBIAS

We employed the TOBIAS (Transcription Factor Occupancy prediction By Investigation of ATAC-seq Signal) (v0.12.10)[34] to identify transcription factor binding sites within peaks detected by ATAC-seq and CUT&Tag-seq. The JASPAR motif database (JASPAR2020_CORE_vertebrates_non-redundant)[35] was used to run TOBIAS with default parameters on narrow peaks identified by MACS2, based on merged biological replicates from both ATAC-seq and CUT&Tag experiments. We then used FOXA1 binding sites identified by TOBIAS to intersect with FOXA1 CUT&Tag peaks, isolating peaks that contain a FOXA1 binding motif. These motif-containing peaks were retained as validated FOXA1 CUT&Tag peaks for subsequent downstream analysis.

### Tissue Sectioning and Visium HD Spatial Gene Expression

5 µm sections were taken from the FFPE tissue blocks with a microtome (HistonCore Biocut). 5 µm sections were taken from the FFPE tissue blocks with a microtome (HistonCore Biocut). Sectioning followed the Visium CytAssist Spatial Gene Expression for FFPE –Tissue Preparation Guide (CG000518, Rev C) for the Visium workflows. We first placed FFPE tissue sections on plain glass slides for deparaffinization, H&E staining and imaging following the Visium HD FFPE Tissue Preparation Handbook (CG000684). Probe hybridization, probe ligation, slide preparation, probe release, extension, library construction, and sequencing followed the Visium HD Spatial Gene Expression Reagent Kits User Guide (CG000685). Sequencing was performed on Illumina Novaseq 6000 with a 150 bp paired-end run.

### Data Analysis for Visium HD

#### Data processing

Data analysis was performed with NovelBrain Cloud Analysis Platform (www.novelbrain.com). We applied fastp with default parameter filtering the adaptor sequence and removed the low quality reads to achieve the clean data. Then we used Space Ranger v3.0.0 to map FASTQ files to the mouse reference, detect the tissue sections, align the sequencing data to the microscope image and the CytAssist image, and output 8×8 µm binned gene-barcode matrices for further analysis.

#### BANKSY Clustering

Sketch based analysis workflows in Seurat (v5.1.0 https://satijalab.org/seurat/) was used for cell normalization and regression based on the expression table according to the UMI counts of each sample to obtain the scaled data. Banksy analysis across multiple samples within Seurat’s spatial framework was performed for joint dimensional reduction and clustering under k_geom=50. Then, harmony-corrected PCA embeddings across samples were constructed on the scaled data with the top 1000 high variable genes, and the top 10 principals were used for UMAP construction. Utilizing graph-based cluster method, we acquired the unsupervised cell cluster result based on the PCA top 10 principal and calculated the marker genes using FindAllMarkers function with wilcox rank sum test algorithm under following criteria:1. log2FC > 0.2; 2. pvalue<0.05; 3. min.pct>0.05.

#### Cell Type Decomposition on Visium HD

We used a published scRNA-seq dataset as a reference, and Visium HD dataset as a spatial query to perform cell annotation with RCTD, a computational approach to deconvolve spatial datasets. And then projected these cell type annotations to the full visium HD spatial dataset.

#### Gene Enrichment Analysis

We applied single cell gene set enrichment analysis based on the public gene sets and normalized gene expression matrix by ssGSEA function in GSVA software package (v1.32.0). And we utilized the R package to draw a boxplot diagram and calculated the difference of the score between the two groups based on Wilcox Rank-sum algorithm.

#### Cell co-occurrence analysis on Visium HD

Cell co-occurrence refers to the occurrence, spatial relationship, of two or more cell types in proximity in the same tissue region. Cell co-occurrence of Stereopy, a fundamental and comprehensive tool for mining and visualization based on spatial transcriptomics data, was calculated based on the cell spatial neighborhood and the distance traverse from 0 to 180 in steps size of 30 with step_num=6 to divide threshold interval evenly. To find the similarities and diversities of cell type cooccurrence among multiple samples, we performed differential co-occurrence among samples.

#### Cell–cell communication analysis

Cell communication analysis was performed using the R package CellChat (v2.0.0) with default parameters. The CellChatDB mouse was used for analysis. The samples were analyzed separately and compared in parallel. This is with the assumption that they were sharing cell types.

#### Differential Gene Expression Analysis

To identify differentially expressed genes among samples, the function FindMarkers with wilcox rank sum test algorithm was used under following criteria:1. Log2FC > 0.25; 2. pvalue<0.05; 3. min.pct>0.1 which was more conducive to obtaining different functional genes among samples.

#### Gene Ontology (GO) analysis

To elucidate the biological implications of the DEGs and marker genes, GO analysis was performed. GO annotations were downloaded from NCBI (http://www.ncbi.nlm.nih.gov/), the Gene Ontology database (http://www.geneontology.org/) and UniProt (http://www.UniProt.org/). Fisher’s exact test was applied to identify the significant GO categories, and an FDR was used to correct the p values.

#### Pathway analysis

Pathway analysis was used to explore the significant pathways of the DEGs and marker genes based on the Kyoto Encyclopedia of Genes and Genomes (KEGG) database. Fisher’s exact test was used to identify significant pathways, and the threshold of significance was defined by the p value and FDR.

#### CNV Estimation

Cells defined as fibroblast were used as reference to identify somatic copy number variations in AT2 cells with the R package infercnv(v0.8.2). We scored each cell for the extent of CNV signal, defined as the mean of squares of CNV values across the genome. Putative malignant cells were then defined as those with CNV signals above 0.05 and CNV correlation above 0.3.

#### Spatial cellular communication networks analysis

For visualizing spatial ligand receptor dynamics in the spatial data, we used the COMMOT package and used the CellChatDB as the ligand-receptor interaction reference database. The ligand–receptor interactions were inferred using Commot’s tl.spatial_communication() function using a distance threshold of 100. Ligand-receptor pair communication directionality was calculated with the tl.communication_direction() function using k□=□5. Vectors for interaction magnitude and directionality were plotted and overlayed onto the spatial image.

### Comparison, Correlation, and Visualization of Gene Expression Data from TCGA and GTEx

Boxplots comparing gene expression between TCGA tumor and normal datasets were generated using TCGA plot[36], an R package designed for integrative pan-cancer analysis and visualization of TCGA multi-omics data. Correlation Scatter Plot of FOXA1 and CDH1 Expression in Normal Lung Tissue Generated from GEPIA[37].

### Ingenuity Pathway Analysis (IPA) Canonical Pathway Analysis

Differentially expressed genes (DEGs) from mRNA-seq data of A549 adenocarcinoma cells, comparing conditions before and after FOXA1 knockdown, were analyzed using Ingenuity Pathway Analysis (IPA). The DEG results were uploaded to the IPA platform as input. For the core analysis, the ‘Expression Analysis’ option was selected as the analysis type, and ‘Expr Log Ratio’ was chosen as the measurement type. Prior to running the analysis, the DEGs were filtered based on the following criteria: a log2 fold change (log2FC) greater than 0.3785, corresponding to a 1.3-fold change, and an adjusted p-value (padj) less than 0.05. Genes that met these thresholds were considered significantly upregulated following FOXA1-KD and were included in the subsequent pathway analysis.

### Western blot

Cell lysates were prepared using RIPA buffer (Thermo, 89900) supplemented with 1% protease inhibitor cocktail (Roche). After a 1-hour incubation on ice, total protein was collected by centrifugation and quantified with a BCA Protein Assay Kit (Beyotime, P0011). Approximately 30 μg of total protein was resolved on a 4–12% SDS–PAGE gel, transferred onto a PVDF membrane (Millipore, ISEQ00010), and blocked with 5% skim milk in TBS–T (Tris-buffered saline containing 0.1% Tween-20) for 1 hour. The membrane was then probed overnight with primary antibodies against FoxA1 (1:1000, CST 53528S), P62 (1:2000, Proteintech 18420-1-AP), cyclin D (1:1000, CST 55506T), mTOR (1:1000, CST 2983T), and GAPDH (1:2000, CST 2118S). The following day, membranes were incubated with HRP-conjugated anti-rabbit IgG (1:200, CST 7074S) and visualized using Chemi-Doc with the BeyoECL Star chemiluminescence kit (Beyotime, P0018AM).

### Immunofluorescence

For immunofluorescence, cells were fixed with 4% paraformaldehyde for 15 min and permeabilized with 0.5% Triton X-100 in PBS for 5 min. After blocking with 3% BSA for 1 h at room temperature, the cells were washed three times in TBS-T (15 min per each wash) and incubated overnight with primary antibodies against lamp1 (1:100, CST 15665S), P63 (1:100, CST 39692S), TFE3 (1:100, Proteintech 14480-1-AP), and CDH1 (1:1000, CST 3195S). Following primary antibodies incubation, the cells were then washed three times in TBS-T (5 min per each wash) and incubated for 1 h in a dark humidity chamber with secondary antibodies Goat Anti-Rabbit IgG H&L (Alexa Fluor® 488) (1:200, ab150077) or Goat Anti-Rabbit IgG H&L (Alexa Fluor® 555), along with Vari Fluor 555-Phalloidin (1:200, MedChemExpress, HY-D1816-300T) and 0.5 mg/mL DAPI. Finally, the cells were washed three times in TBS-T (15 min per each wash) in the dark and mounted onto slide glass using Mowiol solution.

### Cell cycle Assessment

Cell cycle analysis was performed using the Cell Cycle Analysis Kit (no. C1052; Beyotime) according to the manufacturer’s instructions. Cells were harvested and fixed in 70% ethanol for 2 hours at 4°C, then stained with propidium iodide (0.05 mg/ml), RNase A (1 mg/ml), and 0.3% Triton X-100 in the dark for 30 minutes. The distribution of cells in different cell cycle phases was determined by measuring propidium iodide intensity with a BD FACS AriaIII flow cytometer, and the G1, S, and G2/M populations were analyzed using FlowJo VX software.

### Senescence-associated **β**-galactosidase Staining Assay

Cell senescence was assessed using the senescence-associated β-galactosidase Staining Kit (C0602, Beyotime) according to the manufacturer’s instructions. After three washes in PBS, cells were fixed with 4% paraformaldehyde for 15 minutes at room temperature and then incubated overnight at 37°C in the dark with the working solution containing 0.05 mg/mL 5-bromo-4-chloro-3-indolyl-β-D-galactopyranoside (X-gal).

### Organoid Cultures

Single-cell suspensions were prepared from tumor-bearing lung tissues as follows. The tissues were minced under sterile conditions and digested at 37°C for 30 minutes with continuous agitation in Advanced DMEM/F12 medium containing Collagenase Type I (Thermo Fisher Scientific, 450 U/mL) and Collagenase Type IV. The digestion was quenched by adding cold DMEM/F12 supplemented with 10% FBS. The resulting cell suspension was sequentially passed through a 100 μm cell strainer and treated with erythrocyte lysis buffer (eBioscience) to remove debris and red blood cells.

To establish organoid cultures, 5 × 10⁵ tumor cells were resuspended in 250 μL of Matrigel (Corning) and seeded as droplets in 6-well plates. During the first one to two weeks of culture, the Matrigel droplets were overlaid with a recombinant organoid medium. The base medium was Advanced DMEM/F-12, supplemented with 1× B27, 1× N2, 1.25 mM N-acetylcysteine, 100 ng/mL EGF, 125 ng/mL R-spondin1, 100 ng/mL Noggin, 100 ng/mL FGF7, 100 ng/mL FGF10, 0.5 μM SB 2020190, 10 μM Y-27632, GlutaminePlus, HEPES, and Penicillin/Streptomycin (all from Gibco, Sigma, Peprotech, or MCE as specified).

### Transmission Electron Microscope (TEM) Sample Preparation and Visualization

Once cells reached the desired confluency, the culture medium was removed, and 2.5% glutaraldehyde fixative was added at room temperature. Cells were fixed in the dark at room temperature for approximately 5 minutes, then gently scraped in one direction with a cell scraper to avoid repeated scraping. The cell suspension was centrifuged at 2,000 rpm for about 2 minutes, yielding a pellet roughly the size of a mung bean. After discarding the supernatant fixative, the pellet was covered with fresh electron microscopy fixative and left intact. Cells were then further fixed in the dark at room temperature for 2 hours before storage at 4°C.

For lung tissue, fresh samples were cut into cubes of about 2 mm × 1 mm × 1 mm, rinsed in pre-cooled PBS, and immediately immersed in pre-cooled 2.5% glutaraldehyde fixative. After fixation, all samples were rinsed in 0.1 M phosphate buffer and subsequently fixed in 1% osmium tetroxide for 1.5–2 hours. The samples were then washed in 0.1 M phosphate buffer. Following a graded dehydration in ethanol and acetone, the samples underwent resin infiltration and were embedded in pure resin using standard embedding molds. They were polymerized in an electric blast drying oven, and ultrathin sections (50–70 nm) were prepared using a Leica ultramicrotome. The sections were stained with 2% uranyl acetate for 30 minutes, followed by 15 minutes with lead citrate, and then observed with a Hitachi HT7800 TEM at an appropriate magnification.

### Cell Lines and Cell Culture

A549 adenocarcinoma cells (ATCC, Catalog No. CCL-185), RERFLCMS adenocarcinoma cells (Meisen, CTCC-007-0009) and LA759 adenocarcinoma cells (Procell, CL-0376) were cultured in Ham’s F-12K medium (Gibco, #21127030), DMEM (C11960500BT, Gibco), and RPMI 1640 (C11875500CP, Gibco), respectively, with 10% fetal bovine serum (Capricorn scientific, #FBS-52A). All cell culture media were supplemented with penicillin and streptomycin (Gibco, # 15140122). All cells were cultured at 37°C in 5% CO_2_. The 293T cells, kindly provided by Dr. Liu Jing’s laboratory, were maintained in DMEM (C11960500BT, Gibco) containing 10% fetal bovine serum (FBS-52A, Capricorn Scientific) at 37°C in a 5% CO□.

### RNA extraction and RT-qPCR analysis

Total RNA was isolated from cells using the Tissue Total RNA Isolation Kit V2 (Vazyme, #RC112-01). cDNA was synthesized with HiScript III RT SuperMix for qPCR (Vazyme, #R323-01), followed by quantitative PCR (qPCR) performed on the QuantStudio 3 Real-Time PCR System. GAPDH served as the internal control for normalization. The primers were listed in Supplementary Table 1.

### Lyso-tracker Red labeling

Following treatment and washing with warmed complete medium, cells were incubated with 50 nM Lyso-Tracker Red (Beyotime, #C1046) for 1 hour at 37°C to evaluate lysosomal content. Microscopy images were captured immediately following incubation and replacement of the medium with PBS containing Ca²□ and Mg²□. Fields of view were randomly selected and analyzed using ImageJ.

### Transwell Invasion Assay

Approximately 8 × 10□ cells were seeded into the upper compartment of a Transwell plate (Falcon®, #353097) coated with Matrigel basement membrane matrix (Corning, #354234) at a 1:20 dilution. The lower chamber was filled with DMEM supplemented with 10% FBS, and the cells were then cultured at 37°C for 24 hours to allow invasion. Cells that migrated through the Transwell membrane were fixed with 4% formaldehyde, stained with 0.1% crystal violet, and observed under an EVOS microscope (ThermoFisher).

### Vector Design and Lentivirus Generation

The small hairpin RNA (shRNA) sequence of FOXA1 quoted from the GPP Web Portal (#TRCN0000014879, #TRCN0000358450) and the scrambled shNC have been described previously[38], and cloned into a PLKO vector[39].

Exogenous expression of FOXA1 in A549 adenocarcinoma cells was achieved by infecting the cells with a 3×FLAG-FOXA1 expressing lentivirus.

Lentivirus granule was packaged by co-transfecting 293T at 70−80% confluence in a 10cm dish with 4.3 μg of pLKO1, 8.6 μg of psPAX2, and 4.3 μg of pMD2.G using Lipo3000 (Invitrogen, # L3000015). A total of 34 μl of lipo3000 and relevant p3000 reagents per dish were used. After adding the transfection media for 24h, a replacement medium with DMEM containing 20% FBS was used. Lentiviruses were harvested at 72h, and supernatant was centrifuged at 1500 rpm for 10 min to remove cell wreckage and then filtered through a 0.45μm membrane. Adding supernatant to cells, after 24h replaced the culture media.

### Inducible FOXA1-KD Cell Line Construction

The final concentration of doxycycline was 0.5μg/ml maintained 7 days for inducing FOXA1-KD and then adding drugs (with Dox, 0.5μg/ml) 72 hours to generate different EMT states. The final concentrations of drugs TGFβ1 and B1.14 were 10ng/ml and 2μM, respectively.

### Mito Stress Test

The mitochondrial respiration of A549 adenocarcinoma cells bearing lentivirus infection of shNC and shFOXA1 was quantified with an Agilent Seahorse XFe96 Analyzer. 8 × 103 cells were seeded overnight in XF96 V3 PET plates; four peripheral wells contain medium only for background correction. The next day, the XFe96 sensor cartridge was hydrated at 37 °C, CO□-free, first with sterile water and then with 200 µL XF Calibrant for 1 h. Immediately before the run, XF DMEM (pH 7.4) containing 10 mM glucose, 1 mM pyruvate and 2 mM glutamine replaced growth control medium (final volume 180 µL). Oligomycin (1.5 µM), FCCP (2 µM) and rotenone/antimycin A (0.5 µM) were loaded into ports A–C. After calibration, OCR/ECAR were recorded and normalized to cell number.

### Glycolysis Test

Glycolytic activity was quantified with the Seahorse XF Glycolysis Stress Test on an XFe96 Analyzer (Agilent). Briefly, 8 × 10³ per well of A549 adenocarcinoma cells bearing lentivirus infection of shNC and shFOXA1 were seeded in XF96 microplates and incubated overnight in a CO[-free incubator. The next day, growth control medium was replaced with assay medium (XF Base Medium supplemented with 2 mM glutamine, pH 7.4), and the plates were equilibrated for 45 min at 37 °C. Drug stocks (100 mM glucose, 100 µM oligomycin, 500 mM 2-deoxy-D-glucose) were loaded into cartridge ports A–C to achieve final in-well concentrations of 10 mM glucose, 1 µM oligomycin, and 50 mM 2-deoxy-D-glucose; port D remained optional. Extracellular acidification rate (ECAR) data were exported and normalized to cell number.

### Animal Experiments

FOXA1^flox/+^ and *Kras*^G12D^ mice were obtained from Cyagen Co., Ltd (Suzhou, China), while wild-type C57BL/6 mice were purchased from GemPharmatech Co., Ltd (Nanjing, China). All transgenic mice were on a C57BL/6 background. The *Kras*^G12D^, FOXA1^+/+^ and *Kras*^G12D^, FOXA1^flox/+^ strains were generated from the progeny of crosses between FOXA1^flox/+^ and *Kras*^G12D^ mice. All mice were housed under specific pathogen-free conditions. All animal procedures were approved by the Institutional Animal Care and Use Committee (IACUC, No. S2022-508) of Guangzhou Medical University (GMU).

To induce spontaneous lung cancer in the mouse model, control group (*Kras*^G12D^, FOXA1^+/+^) and FOXA1-KD (*Kras*^G12D^, FOXA1^flox/+^) mice were administered Sftpc-Cre-expressing AAV9 particles (5 × 10¹¹ viral genomes in 200 µL PBS) via tail-vein injection. After 30 days, the lungs were harvested for subsequent analyses.

To generate cell line tail vein injection into NCG mice (obtained from GemPharmatech Co., Ltd) lung for evaluating tumorigenesis of lung cancer cell lines, 1×10^6^ A549 adenocarcinoma cells were suspended in 200 μL of PBS (Gibco) and gently injected into the mouse tail vein using a 1 mL sterile syringe (Biosharp).

Before the injection of doxycycline-induced KD or control cells, NCG mice received 2 mg/mL doxycycline orally once daily, with water changed every 2 days, for a total of 14 days before tail vein injection. After injection, doxycycline administration continued until the mice were sacrificed.

To evaluate the tumorigenic potential of lung cancer cell lines, 1×10^6^ A549 adenocarcinoma cells were suspended in 200 μL of PBS (Gibco) and gently injected into the tail vein of NCG mice using a sterile 1 mL syringe (Biosharp). Before the injection of doxycycline-induced knockdown or control cells, NCG mice received 2 mg/mL doxycycline orally once daily, with the drinking water changed every 2 days, for a total of 14 days prior to tail vein injection. After injection, doxycycline administration continued until the mice were sacrificed.

### shRNA-expressing Stable Cell Screening

For Tet-inducible FOXA1 shRNA-expressing stable cells screening, FOXA1 shRNA oligomers were synthesized (sequences are shown in Supplementary Table 1), annealed, and cloned into a Tet-pLKO-puro cloning vector (Addgene #47541) at AgeI and EcoRI sites. FOXA1 shRNAs were introduced into LA795 cells by lentiviral infection, and the infected cells were subsequently screened with Puromycin for 5-7 days at a final concentration of 3 3μg/ml. The remaining cells, which integrated the shRNA-expressing cassette, were used as a cell pool to detect FOXA1 expression and perform other functional tests. For constitutive FOXA1 shRNA-expressing stable cells screening, FOXA1 shRNA oligomers were synthesized (sequences are shown in Supplementary Table 1) annealed, and cloned into a pGIPZ-miR-30b-3p cloning vector (P10719, MiaoLingBio) at XhoI and EcoRI sites[40] to construct vectors; all subsequent steps were the same as described above.

## Results

### A new TGF-β receptor inhibitor prevents EMT in A549 lung adenocarcinoma cells and increases tumorigenicity

The A549 non-small cell lung adenocarcinoma (NSCLC) cell line exhibits an intermediate plastic EMT phenotype, which can be induced to a more mesenchymal phenotype with TGF-β or an epithelial-like phenotype with (TGF-βR) inhibitors [41, 42]. We employed the TGF-β receptor inhibitor A83-01 to synthesize compound B1.14, which exhibited selective and effective binding affinity toward the TGF-β receptor type I (TGFBR1) (Supplementary material 1 and Extended Fig. 1 A-E). A549 adenocarcinoma cells were treated with TGF-β, DMSO, or the small-molecular-weight compound TGF-βR inhibitor B1.14 together with TGF-β for 72 hours to identify key factors positioning these cells along the EMT spectrum. Cells exposed to TGF-β displayed morphological changes characteristic of EMT, appearing elongated compared to the DMSO-treated control group (Fig. 1A). In contrast, A549 adenocarcinoma cells treated with B1.14 in combination with TGF-β (hereafter referred to as B1.14) exhibited an epithelial phenotype, characterized by the formation of compact, tightly clustered cell colonies (Fig. 1A). B1.14 demonstrated superior efficacy compared with the TGF-β inhibitors A83-01 and Galunisertib in suppressing invasion and migration of A549 lung adenocarcinoma cells (Extended Fig. 2A–D). RNA sequencing analysis, conducted after 72 hours of treatment, revealed regulatory proteins, along with morphological changes (Fig. 1B). EMT scoring[43] and Gene set enrichment analysis (GSEA) demonstrated that B1.14, DMSO, and TGF-β treatments produced epithelial, intermediate, and mesenchymal EMT signatures, respectively (Fig. 1C,D, Extended Fig. 2E).

**Fig. 1.**
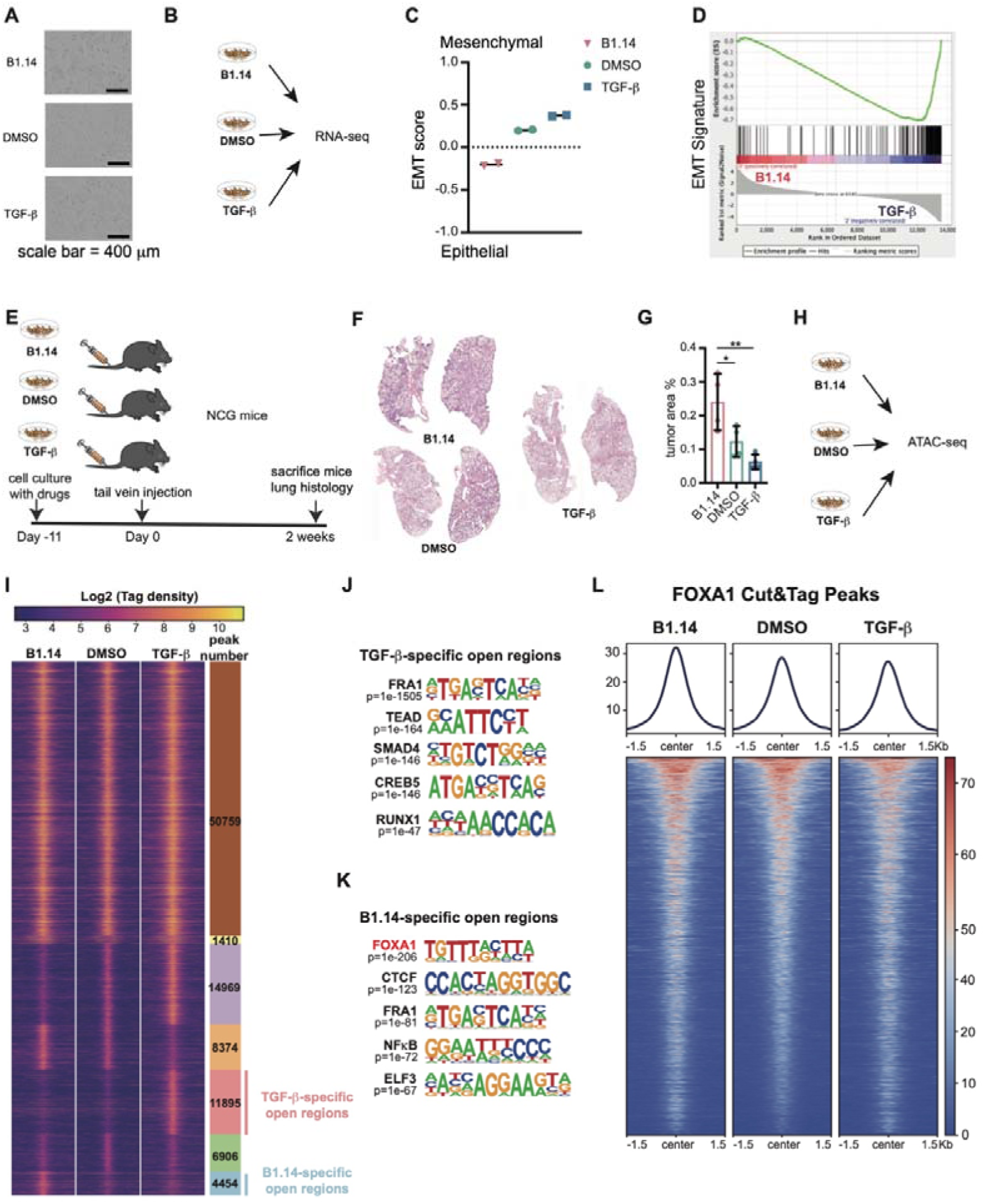
FOXA1 exhibits strong binding in the epithelial-like state of lung adenocarcinoma cells. **A**, A549 adenocarcinoma cells treated with B1.14, DMSO, and TGF-β, respectively. **B**, RNA-seq workflow on cells treated with B1.14, DMSO, and TGF-β, respectively. **C**, EMT scores derived from RNA-seq data are analyzed for A549 adenocarcinoma cells treated with B1.14, DMSO, and TGF-β, respectively. **D**, Gene Set Enrichment Analysis (GSEA) of the Hallmark EMT pathway is performed to compare A549 adenocarcinoma cells treated with B1.14 to those treated with TGF-β. **E**, A549 adenocarcinoma cells treated with B1.14, DMSO, or TGF-β were injected into the tail veins of NCG mice. Two weeks post-injection, tumor formation is assessed by histological examination of the left lung sections. **F**, Representative H&E staining of lung sections from each experimental group are presented for comparative analysis. **G**, A bar plot displays the percentage of tumor area, quantified using HALO software. Statistical analysis is conducted using one-way ANOVA. **H**, ATAC-seq workflow is conducted on cells treated with B1.14, DMSO, and TGF-β, respectively. **I**, ATAC-seq peak profiles for A549 adenocarcinoma cells treated with B1.14, DMSO, and TGF-β, respectively. **J**, The top five transcription factor motifs identified in TGF-β−specific open chromatin regions. **K**, The top five transcription factor motifs identified in B1.14-specific open chromatin regions. **L**, FOXA1 Cut&Tag peak profiles obtained for A549 adenocarcinoma cells treated with B1.14, DMSO, and TGF-β, respectively.

To compare the tumorigenic potential of A549 adenocarcinoma cells at distinct stages of EMT, we injected these cells with either an epithelial phenotype (B1.14) or a mesenchymal phenotype induced by TGF-β into the tail veins of NCG mice lacking B, T, and NK cells (Fig. 1E). After 2 weeks of tumorigenesis in NCG mice, epithelial-like cells generated markedly larger pulmonary tumors (Fig. 1F,G) than their mesenchymal counterparts (Fig. 1F,G). These data demonstrated that the EMT status critically influences lung tumor formation in vivo, highlighting the importance of dissecting the mechanisms governing EMT in NSCLC.

### FOXA1 exhibits heightened chromatin occupancy in epithelial-like A549 adenocarcinoma cells

To further explore regulatory factors driving morphological transition in A549 adenocarcinoma cells, we categorized ATAC-seq data based on chromatin accessibility in epithelial (B1.14-treated), intermediate (DMSO-treated), and mesenchymal (TGF-β-treated) conditions (Fig. 1H). A549 adenocarcinoma cells acquired different regulatory patterns in these three EMT states as reported previously[44] (Fig. 1I). TGF-β increased chromatin accessibility more than in the TGF-β receptor inhibitor-treated cells. Motif analysis using the HOMER de novo method identified transcription factors enriched in mesenchymal (Fig. 1J) and epithelial (Fig. 1K) states. In TGF-β-treated mesenchymal-like A549 adenocarcinoma cells, FRA1, TEAD, SMAD4, CREB5, and RUNX1 motifs were identified as the most significantly accessible genes. These transcription factors were previously found to be associated with various carcinoma cells undergoing EMT[45–49]. FOXA1, CTCF, FRA1, NF-κB, and ELF3 motifs were enriched in B1.14 epithelial-like A549 adenocarcinoma cells. FOXA1 was the most enriched DNA-binding protein in the epithelial state of A549 adenocarcinoma cells, suggesting its potential role in sustaining the epithelial phenotype. CTCF is also associated with an epithelial phenotype, as its downregulation by miR-137 promoted EMT in esophageal squamous cell carcinoma[50]. FRA1 is considered a transcription factor inducing EMT-related genes[51, 52], possibly due to some residual TGF-β signaling rendering accessible the FRA1 motif in B1.14 and TGF-β treated group. NF-κB signaling pathway was associated with EMT in lung cancer development[53]. The mRNA level of NF-κB2, a major transcription factor in non-canonical NF-κB signaling, was significantly elevated in B1.14-treated epithelial-like A549 adenocarcinoma cells (Extended Data Fig. 2F). As for FRA1, TGF-β residual signaling may have induced NF-κB in B1.14-treated cells without altering the epithelial phenotype. Inhibition of the ELF3 transcription factor induces EMT through TGF-β signaling in A549 and PC9 adenocarcinoma cells[54], supporting a role in maintaining the epithelial state, as shown in our ATAC-seq analysis.

We then focused on FOXA1, which is associated with an epithelial phenotype in the development and differentiation of the airway epithelium[55] as well as essential for reprogramming the epigenetic landscape in mouse *Kras* mutation lung adenocarcinoma models[12]. To elucidate FOXA1 binding dynamics across the epithelial–mesenchymal transition (EMT) spectrum in lung adenocarcinoma, FOXA1 Cut&Tag sequencing was conducted on A549 adenocarcinoma cells subjected to treatment with B1.14, DMSO, or TGF-β. The results confirmed significant FOXA1 binding in the epithelial state (Fig. 1L). Analysis of the TCGA LUAD dataset further revealed that FOXA1 expression was higher in tumor samples than in normal lung tissue, suggesting a role for FOXA1 in the tumorigenesis of LUAD (Extended Data Fig. 2G).

As previously shown, A549 adenocarcinoma cells treated with B1.14, in which FOXA1 binding was highly enriched in chromatin-accessible regions, formed larger tumors (Fig. 1F,G) than TGF-β-treated cells, in which FOXA1 binding was minimal (Fig. 1F,G). These findings highlight FOXA1 as a critical regulator of the epithelial state associated with robust proliferation.

### FOXA1 is highly expressed in polarized and proliferating carcinoma cells in the Kras murine model

To investigate FOXA1 expression in mouse adenocarcinoma, a *Kras*^G12D^ mouse model was established by introducing an inducible *Kras* mutation in AT2 cells[56]. Primary LUAD tumors were induced by tail vein injection of an AAV9 vector encoding Cre recombinase driven by a Sftpc-specific promoter to induce solely the *Kras*^G12D^ mutation in AT2 cells (Fig. 2A). Mice developed substantial lung tumors at 90 days following *Kras* mutation activation in AT2 cells.

**Fig. 2.**
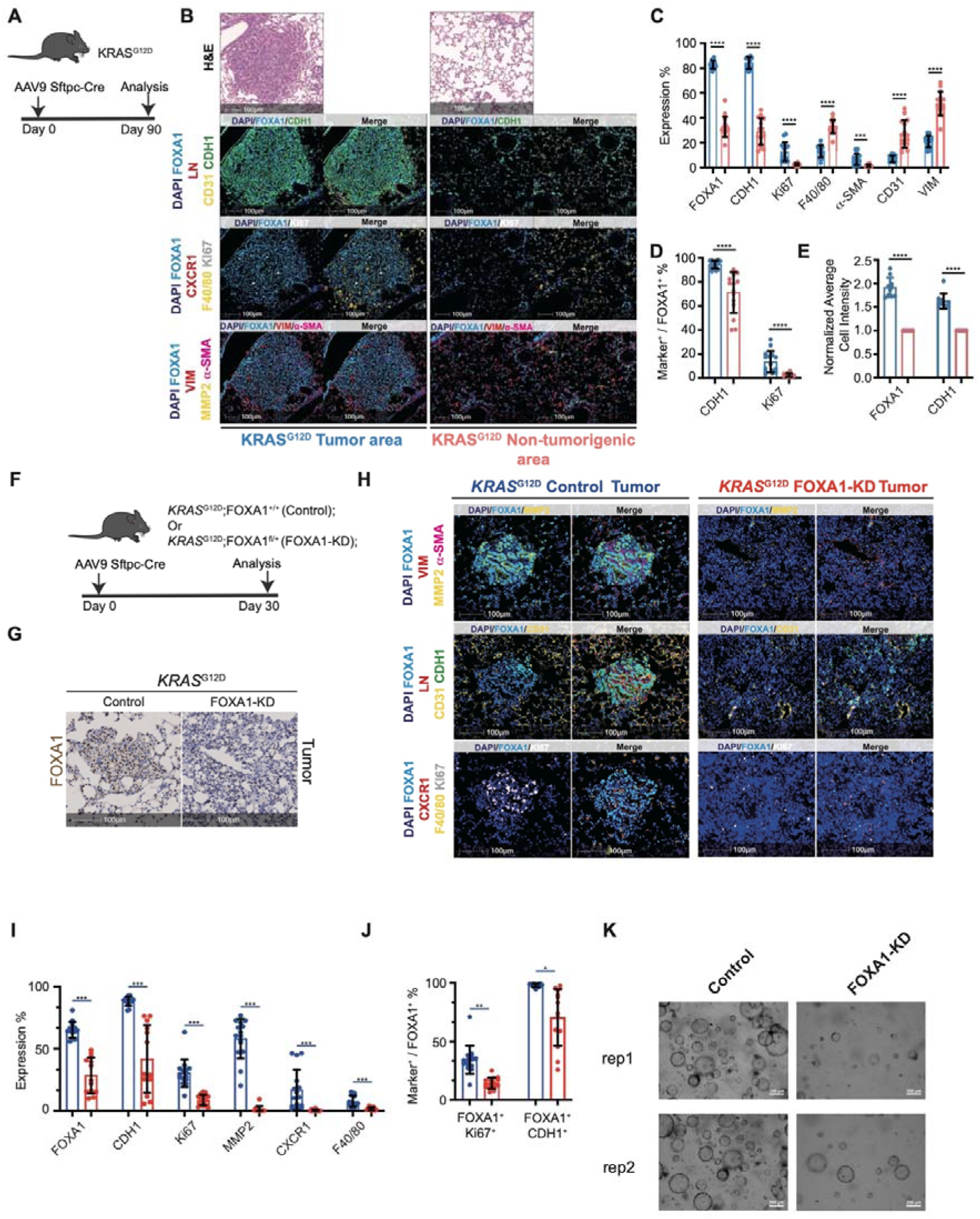
FOXA1 is highly expressed in *Kras*-driven polarized adenocarcinoma cells and promotes tumorigenic. **A**, Scheme for generating *Kras*^G12D^ lung adenocarcinoma. **B**, H&E and mIHC staining on serial sections from tumor and non-tumor regions. **C**, The proportion of cells expressing positive markers is calculated as a fraction of the total cell population, and statistical significance was assessed using one-way ANOVA. **D**, The proportion of cells double-positive for FOXA1, CDH1, or KI67 is calculated among all FOXA1-positive cells, and statistical analysis was performed using one-way ANOVA. **E**, Normalized average cellular intensities of FOXA1 and CDH1 are quantified, and statistical analysis is conducted using one-way ANOVA. **F**, Scheme for generating *Kras*^G12D^ FOXA1^+/+^ and *Kras*^G12D^ FOXA1^fl/+^ lung adenocarcinoma models. **G**, IHC staining for FOXA1 on tumor regions from *Kras*^G12D^ FOXA1^+/+^ and *Kras*^G12D^ FOXA1^fl/+^ samples. **H**, MIHC staining on serial sections from *Kras*^G12D^ FOXA1^+/+^ and *Kras*^G12D^ FOXA1^fl/+^ tumor regions. **I**, The percentage of marker-positive cells is calculated relative to the total cell population, and statistical significance was determined using one-way ANOVA. **J**, The proportion of cells co-expressing FOXA1, CDH1 or KI67 is calculated among FOXA1-positive cells, with statistical analysis conducted using one-way ANOVA. **K**, Tumor organoids are generated separately from *Kras*^G12D^ FOXA1^+/+^ and *Kras*^G12D^ FOXA1^fl/+^ tumors. scale bar = 20 μm.

Multiplex immunohistochemistry (mIHC) staining on paraffin sections of mouse LUAD was performed to establish the distribution of FOXA1, CDH1, VIM, and LN as epithelial cell polarity and EMT-related markers, as well as CD31, CXCR1, MMP2, α-SMA, and F4/80 as TME-related markers, and Ki67 as a proliferation marker.

These analyses revealed that FOXA1 was highly overexpressed in carcinoma cells compared to normal lung epithelial AT2 cells (Fig. 2B). FOXA1 was predominantly expressed in the nucleus of *Kras*-induced LUAD cells, which formed apical-basal polarized structures with CDH1 expressed at cell-cell junctions (Fig. 2B). In contrast, the non-tumorigenic regions contained large numbers of macrophages and stromal cells, many of which were VIM-positive, while fewer stromal cells were found within the tumor bed.

Interestingly, most macrophages in the tumor periphery were CXCR1^+^ but were absent within the tumor bed (Fig. 2B). CXCR1^+^ macrophages, are considered senescent-like in *Kras*-driven lung cancer progression[57]. FOXA1-positive carcinoma cells exhibited higher Ki67 staining than peripheral regions where only proliferating AT2 cells could be positively stained (Fig. 2B), supporting tumor growth associated with the epithelial tumor state.

LN is expressed at the basal surface of the polarized carcinoma cell islands[58]. In the non-tumorigenic region, the mIHC results showed LN co-localized with CD31, a vascular endothelial marker[59]. We observed that LN was present on the basal side of carcinoma cells, regardless of the presence of endothelial cells within the tumor bed (Fig. 2B). LN contributes to sustaining the apical-basal polarity of carcinoma cells, rather than being solely associated with the endothelium.

To quantify marker expression levels in mIHC staining, statistical analyses were performed using the HALO software to measure the cell number and intensity per cell associated with each marker and their co-localizations. Tumor regions showed significantly higher expression of FOXA1, CDH1, Ki67, and α-SMA compared to non-tumorigenic areas, while CD31, VIM, and F40/80 were more abundantly expressed in non-tumorigenic regions (Fig. 2C). Notably, nearly all FOXA1-positive LUAD cells expressed CDH1, compared to 70% in non-tumor areas containing AT2, AT1 and some other cells like endothelial and stromal cells (Fig. 2D). The expression intensity of FOXA1 and CDH1 was also elevated in carcinoma cells (Fig. 2E).

Furthermore, the co-expression of FOXA1 and Ki67 was dominant in these regions, highlighting FOXA1-positive carcinoma cells as key contributors to tumor initiation and progression in *Kras*^G12D^-driven LUAD (Fig. 2D).

### Attenuated FOXA1 expression in Kras-driven tumors alters polarization and decreases tumorigenicity

*Kras*^G12D^; FOXA1^fl/+^ (hereafter referred to as FOXA1-KD) mice were generated by crossing *Kras*^G12D^ mice with FOXA1^fl/+^ mice. This model was established to investigate whether reduced FOXA1 impacts lung adenocarcinoma cell polarization as well as primary lung carcinoma development in parallel with *Kras*^G12D^; FOXA1^+/+^ mice (hereafter referred to as control mice)[60] (Fig. 2F). Mice were euthanized 30 days following induction of Sftpc-Cre, during which a subset of the infected AT2 cells exhibited overexpression of the *Kras*^G12D^ protein, thereby promoting the development of lung adenocarcinoma. A significant decrease in FOXA1 expression was observed in the lung sections of Cre-induced FOXA1-KD mice through FOXA1 IHC staining (Fig. 2G).

Transmission electron microscopy (TEM) was performed to further characterize the phenotype of lung adenocarcinoma cells in control and FOXA1-KD *Kras*^G12D^ mice that had been induced with Sftpc-Cre for 30 days. Abundant polarized LUAD cells were observed in the tumor bed of control *Kras*^G12D^ mice (Extended Data Fig. 3A). In contrast, carcinoma cells in FOXA1-KD mice lacked polarized structures and exhibited smaller intercellular junctions in FOXA1-KD *Kras*^G12D^ adenocarcinoma cells (Extended Data Fig. 3A).

Ki67 expression was markedly elevated in regions with abundant FOXA1 levels (Fig. 2H,I), and cells co-expressing FOXA1 and Ki67 were significantly enriched in control tumors (Fig. 2J).

To more precisely assess the tumorigenic potential, three-dimensional tumor organoid cultures were established from enzymatically dissociated neoplastic lesions from control and FOXA1-KD *Kras*^G12D^ mice[61]. Adenocarcinoma cells with diminished FOXA1 expression from FOXA1-KD tumors generated fewer organoids than in control groups (Fig. 2K). Given that lung cancer organoids recapitulate cellular plasticity, stem-like properties, and robust proliferation[62], this loss of organoid-forming capacity indicates that FOXA1 downregulation impairs tumorigenicity.

### K*ras*-driven adenocarcinoma cells with reduced FOXA1 expression accumulate lysosome-like organelles

TEM of lung adenocarcinoma cells revealed not only a reduction in cell–cell junctions in the FOXA1-KD *Kras*^G12D^ mice, but also alterations in cell morphology and organelle composition.

We noticed that the nuclear morphology of proliferating adenocarcinoma cells differed between control and FOXA1-KD *Kras*^G12D^ mice (Extended Data Fig. 3A). Nuclei exhibited irregular contours and lobulations in adenocarcinoma cells in FOXA1-KD mice, in contrast to the more rounded nuclei of adenocarcinoma cells in control mice. Nuclear dysmorphia is a significant attribute of accentuated cancer malignancy[63–65].

Interestingly, we also observed that the cytoplasm of adenocarcinoma cells in FOXA1-KD mice was enriched with large lysosome-like organelles (Extended Data Fig. 3A). This observation suggested that the accumulation of lysosome-like organelles in adenocarcinoma cells may result from the reduced FOXA1 expression.

### K*ras*-driven lung adenocarcinoma foci with reduced FOXA1 expression exhibit an altered TME

MIHC staining was performed for FOXA1, CDH1, VIM, CD31, F40/80, CXCR1, α-SMA, MMP2, LN, and Ki67 in control and the FOXA1-KD *Kras*^G12D^ tumor regions.

Through quantitative measurement of fluorescence-labeled cells, FOXA1-KD tumors showed a decrease in CDH1-positive cells, indicating a loss of polarity that affects the epithelial phenotype in tumors with reduced FOXA1 expression (Fig. 2I). FOXA1 and CDH1 double-positive cells were also lost as FOXA1 expression decreased along with Ki67 (Fig. 2J).

In addition to Ki67 and CDH1, the control tumor regions contained significantly more MMP2 and CXCR1-positive cells than the FOXA1-KD regions, and control TME was notably enriched with macrophages (Fig. 2H,I). A MMP2-enriched microenvironment has been linked to the early development of non-small cell lung cancer in human patients[66], suggesting that the potential for lung cancer progression is greater in control tumors compared to FOXA1-KD tumors (Fig. 2I). CXCR1-positive macrophages were also found to be enriched in control lung cancer regions (Fig. 2H,I). These CXCR1^+^ macrophages are considered to exhibit a senescence-like phenotype and are thought to promote tumorigenic progression by modulating the local TME[57].

Collectively, these findings suggest that reduced FOXA1 expression in *Kras*^G12D^-driven murine adenocarcinoma cells reshapes the TME, which may underline the phenotypic changes observed in the context of FOXA1 deficiency.

### *Kras*-driven adenocarcinoma cells deficient in FOXA1 exhibit increased malignancy

We observed that lung adenocarcinoma cells with reduced FOXA1 expression exhibited decreased polarity, an accumulation of lysosome-like organelles, and were associated with alterations in the TME. To gain further insight into the mechanisms driving the observed modifications in lung adenocarcinoma cells following *Kras*^G12D^ FOXA1-KD, HD spatial sequencing was performed on lung tissue sections from control *KrasG*^12D^ mice and FOXA1-KD *Kras*^G12D^ mice after 30 days of Cre recombinase induction (Fig. 3A). Following single-cell RNA-seq analysis of a *Kras*^G12D^ lung adenocarcinoma[67], a total of 318,910 cells in control mice and 296,845 cells in FOXA1-KD mice were successfully annotated (Fig 3A, Extended Data Fig. 4A-G, Extended Data Fig. 5A-D).

**Fig. 3.**
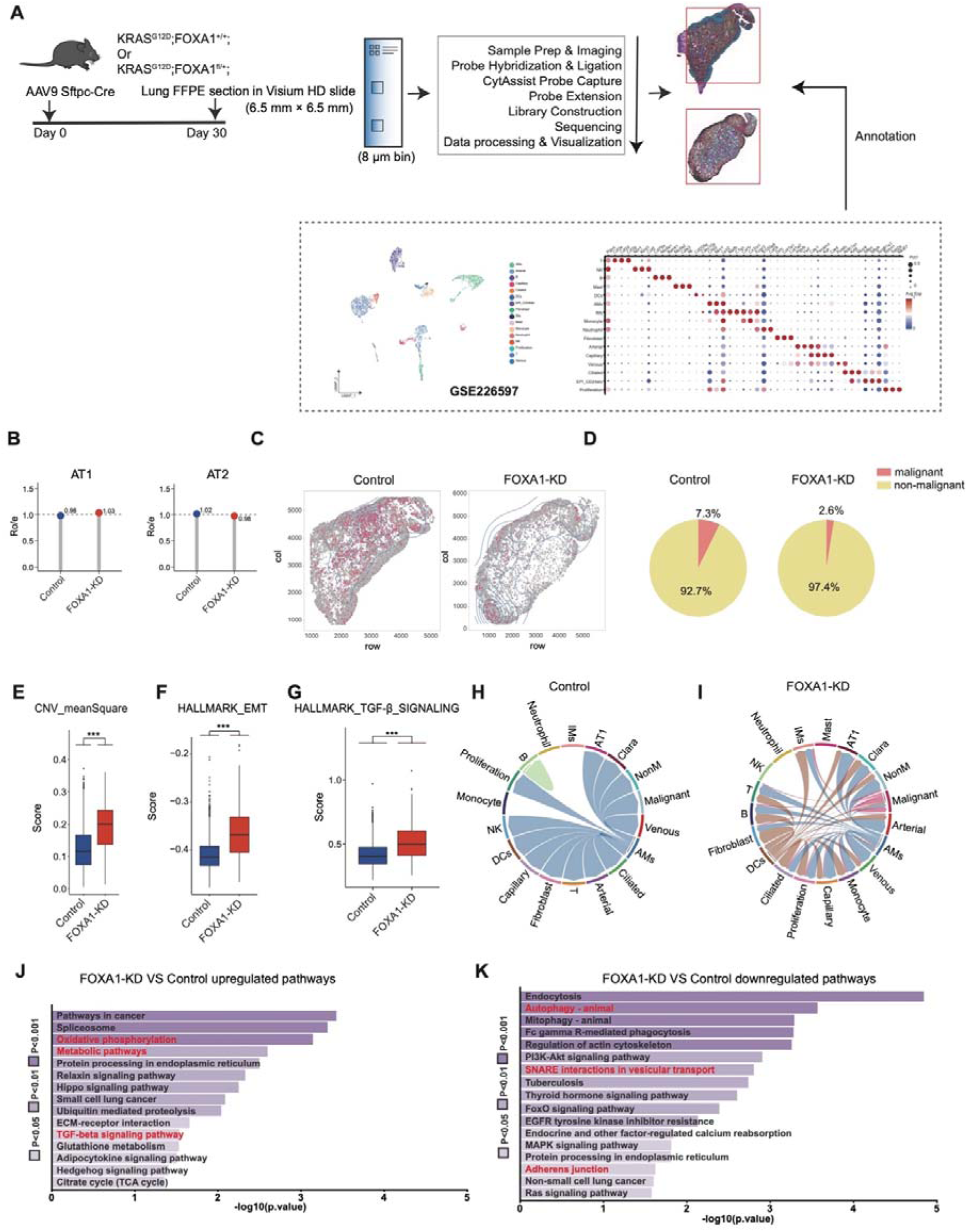
FOXA1-KD *Kras*-driven tumors exhibit enhanced malignancy through elevation of TGF-β signaling pathway. **A**, Workflow of Visium HD spatial analysis and annotation in control and FOXA1-KD mouse tumors. **B**, Ro/e of cell number comparison of AT1 and AT2 in control and FOXA1-KD mouse tumors. **C**, Distribution of malignant cells in control and FOXA1-KD K*ras*^G12D^ samples. **D**, Proportion of malignant cells among all AT2 cells in control and FOXA1-KD K*ras*^G12D^ samples. **E**, Higher CNV meanSquare value indicates higher malignancy. FOXA1-KD malignant cells show higher abnormal score compared to control malignant cells. **F**, HALLMARK EMT score calculated for malignant cells in control and FOXA1-KD K*ras*^G12D^ samples. **G**, HALLMARK TGF-β signaling score calculated for malignant cells in control and FOXA1-KD K*ras*^G12D^ samples. **H**, Chord diagrams show that TGF-β signaling crosstalk among the microenvironment cells in control K*ras*^G12D^ samples. **I**, Chord diagrams show that TGF-β signaling crosstalk among the microenvironment cells in FOXA1-KD K*ras*^G12D^ samples. **J**, Upregulated pathways in FOXA1-KD adenocarcinoma cells compare to control cells. **K**, Downregulated pathways in FOXA1-KD malignant cells compare to control malignant cells.

A total of 118,085 alveolar type II (AT2) and 58,435 alveolar type I (AT1) cells were identified in the control mice, and 76,083 AT2 and 41,476 AT1 cells were found in FOXA1-KD mice (Extended Data Fig. 5E-G). The observed to expected cell number ratio (Ro/e) was calculated for AT1 and AT2 populations in both control and FOXA1-KD mice[68]. AT1 cells were preferentially enriched in the FOXA1-KD mice, whereas AT2 cells predominated in control mice (Fig. 3B, Extended Data Fig. 5E-G).

The selective activation of the *Kras*^G12D^ allele in alveolar type II (AT2) cells markedly promoted aberrant AT2 proliferation, consequently expanding the AT2 cell population. InferCNV (https://github.com/broadinstitute/inferCNV) was applied to delineate malignant clones by evaluating chromosomal copy number variation (CNV) within the AT2 compartment and quantitatively assessing their oncogenic status (Extended Data Fig. 6A,B). The spatial distribution of malignant cells (Fig. 3C) paralleled FOXA1 immunohistochemical staining in control and FOXA1-KD sections (Extended Data Fig. 6C). FOXA1-KD samples showed diminished staining intensity and fewer FOXA1-positive cells relative to controls. InferCNV not only discriminates malignant from normal cells but also quantifies chromosomal instability through CNV scores[69]. Although the FOXA1-KD mice contained a smaller fraction of malignant cells (2.6% versus 7.3% malignant cells in control mice (Fig. 3D), these cells exhibited a higher degree of malignancy than those in the control group (Fig. 3E).

High-definition spatial sequencing revealed that reducing FOXA1 expression in AT2 cells decreased the number of transformed cells, and these cells displayed greater chromosomal instability making them more malignant than their counterparts in control mice.

### FOXA1-KD induces a mesenchymal phenotype characterized by elevating TGF-β signaling and lysosomes through reduced SNARE expression

MIHC staining showed that lung adenocarcinoma cells in the FOXA1-KD tumors expressed lower levels of CDH1 (Fig. 2J). Hallmark EMT scores calculated from high-definition spatial sequencing of malignant cells showed that adenocarcinoma cells in the FOXA1-KD tumors displayed a more pronounced EMT phenotype than those in the FOXA1-control tumors (Fig. 3F). These results confirmed that FOXA1 was a key regulator of the epithelial state in lung adenocarcinoma cells.

In NSCLC, EMT is closely associated with malignancy, promoting invasion, metastasis, and drug resistance[70, 71]. TGF-β signaling is a principal inducer of EMT in NSCLC and drives tumor progression[42]. Notably, FOXA1 exhibited increased binding activity upon inhibition of the TGF-β signaling pathway (Fig. 1L). To assess whether FOXA1 depletion enhances TGF-β signaling, pathway activity scores were calculated in malignant cells from control and FOXA1-KD mice. TGF-β signaling was significantly up-regulated in FOXA1-KD adenocarcinoma cells, indicating that FOXA1 inhibits TGF-β signaling (Fig. 3G).

Elevated TGF-β signaling is thought to be associated with alterations in the TME[42, 72]. Cell–cell communication analysis of the TGF-β pathway indicated a markedly more complex signaling landscape in the FOXA1-KD tumors. TGF-β produced by the malignant cells paracrinally activates multiple cell types in the TME, whereas no such signaling pathway was detected in the control tumors (Fig. 3H,I). Cell co-occurrence analysis further revealed that malignant cells in the FOXA1-KD tumors co-localized with a broader array of cell types in the TME than those in the control tumors (Extended Data Fig. 6D,E).

Pathway enrichment analysis of the FOXA1-KD and control tumors (Fig. 3J,K) revealed marked differences in their molecular programs. FOXA1-KD tumors, in addition to enhanced TGF-β signaling, exhibited widespread upregulation of metabolic pathways and a preferential enrichment of gene sets characteristic of small-cell lung cancer. We also observed that control tumors showed heightened activity of the adherens junction pathway, consistent with more pronounced cell polarity.

Since the pathway analysis did not reveal an upregulation of lysosome biogenesis in FOXA1-KD *Kras*^G12D^ malignant cells, we inferred that the observed lysosomal accumulation in FOXA1-KD *Kras*^G12D^ adenocarcinoma cells (Extended Data Fig. 3A) occurs independently of lysosome biogenesis pathways. In contrast, compared to the control tumors, FOXA1-KD *Kras*^G12D^ malignant cells showed a significant reduction in the “SNARE interactions in vesicular transport” pathway. Soluble N-ethylmaleimide-sensitive factor attachment protein receptor (SNARE) proteins family is essential for lysosomal function and the regulation of intracellular membrane fusion, with their dysfunction strongly associated with an aberrant lysosomal accumulation. Loss of SNARE family members such as SNAP25 and VAMP8 disrupts the autolysosomal flux, leading to the accumulation of autophagic substrates in lung cancer cells, highlighting the critical role of SNAREs in lysosomal homeostasis[73]. In hepatocellular carcinoma cells, impaired assembly of the STX17-SNAP29-VAMP8 complex hampers autophagosome-lysosome fusion and promoting abnormal lysosome accumulation[74]. Similarly, SNAP29 deficiency restricts autophagosome–lysosome fusion, resulting in the accumulation of autophagic substrates and disrupting cell homeostasis[75]. Moreover, control tumors were enriched for mitophagy and autophagy signatures, suggesting that although lysosomes accumulated in FOXA1-KD tumors, their functionality appeared to be reduced compared with that in control tumors.

### FOXA1 is overexpressed in well-differentiated epithelial-like human LUAD and is associated with TME remodeling

We next examined whether FOXA1 expression in human LUAD parallels the pattern observed in *Kras*^G12D^-driven murine LUAD. Human LUAD is commonly characterized by TTF1-positive labeling[76]. We stained TTF1 in well-differentiated lepidic-predominant LUAD patient tissue sections (n=5). To elucidate the spatial relationship between FOXA1 and TTF1 in human lung adenocarcinoma tissue, and to assess the cellular characteristics of FOXA1-positive cells, we conducted mIHC staining for FOXA1, VIM, α-SMA, and CDH1 on serial sections adjacent to those subjected to TTF1 immunohistochemistry (Fig. 4A). TTF1 and FOXA1 colocalized within the same adenocarcinoma and AT2 cells (Fig. 4A). Previous studies using mouse and human LUAD samples have demonstrated that FOXA1 and FOXA2 can cooperatively regulate TTF1 expression in adenocarcinoma cells[12], supporting the association between FOXA1 and TTF1.

**Fig. 4.**
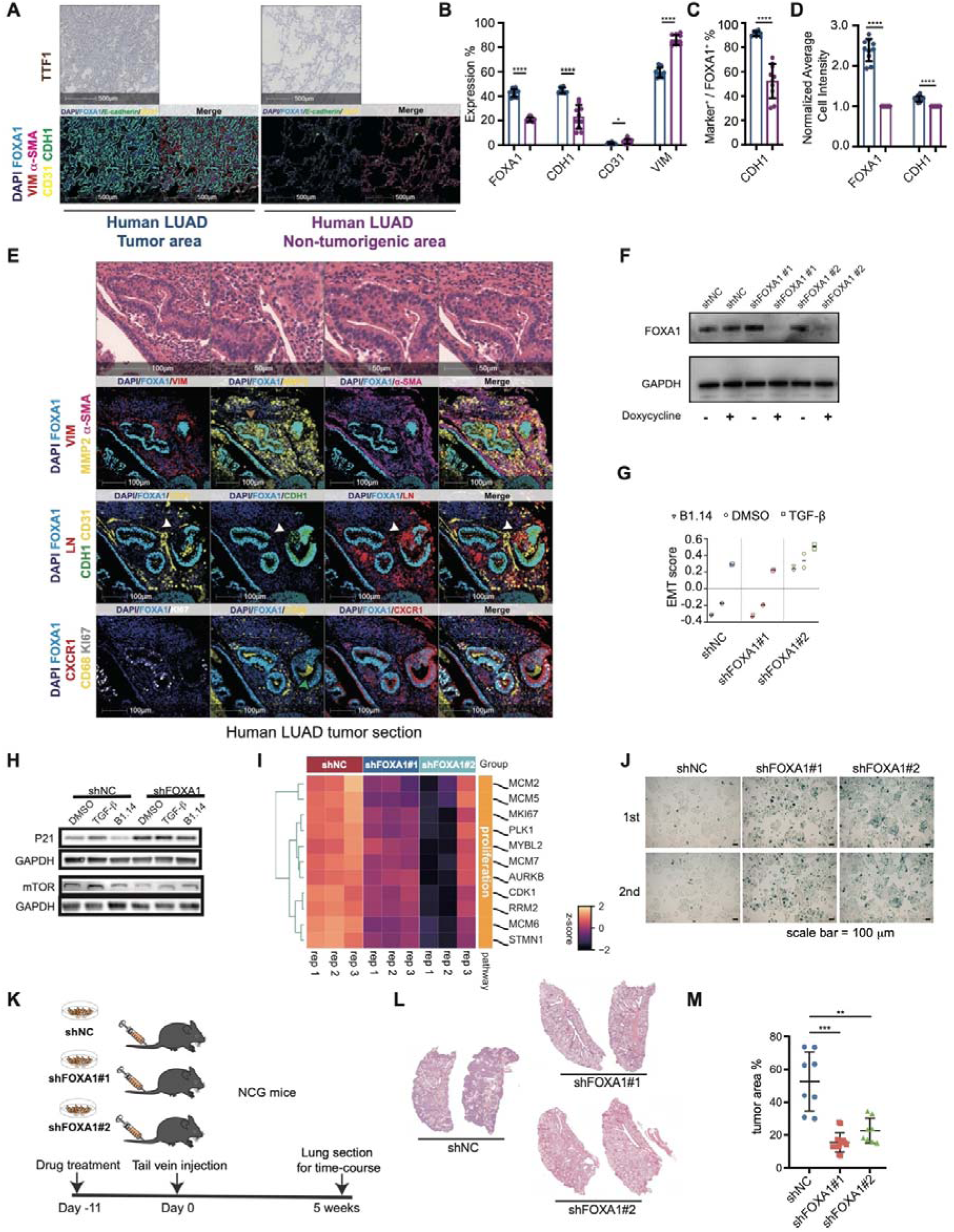
FOXA1 is highly expressed in human polarized LUAD samples and facilitates cellular proliferation as well as tumorigenic capacity. **A**, Immunohistochemical TTF1 staining and mIHC staining on adjacent sections of human LUAD specimens encompassing both tumor and non-tumor regions. **B**, The proportion of cells expressing positive markers is calculated as a fraction of the total cell population, and statistical significance was assessed using one-way ANOVA. **C**, The proportion of cells double-positive for FOXA1 and CDH1 is calculated among all FOXA1-positive cells, and statistical analysis was performed using one-way ANOVA. **D**, Normalized average cellular intensities of FOXA1, and CDH1, statistical analysis is conducted using one-way ANOVA. **E**, H&E and mIHC staining of human LUAD tissue serial sections obtained from both tumor regions. Brown arrow indicates carcinoma cells with low FOXA1 expression and high MMP2 expression; white arrows indicate regions with low FOXA1 expression and lacking polarization. **F**, Western blot analysis of FOXA1 in FOXA1-KD A549 adenocarcinoma cells. **G**, EMT score of shNC and shFOXA1#1 and shFOXA1#2 A549 adenocarcinoma cells. **H**, Western blot analysis of p21 and mTOR in shNC- and shFOXA1#1-transduced cells following treatment with B1.14, DMSO, or TGF-β. **I**, Gene expression profiles of proliferation-related genes in shNC and shFOXA1 cells. **J**, β-galactosidase staining of shNC, shFOXA1#1 and shFOXA1#2 A549 adenocarcinoma cells. **K**, Experimental workflow: shNC- or shFOXA1-transduced cells are injected via the tail vein of NCG mice to establish lung tumors. **L**, Representative histological lung sections obtained 5 weeks after cell injection. **M**, Dot-plot summarizing the quantitative analysis of tumor area in lung sections 5 weeks after tail vein injection. Statistical analysis is performed using one-way ANOVA.

FOXA1, VIM, α-SMA, and CDH1 exhibited comparable expression profiles in both human and murine LUAD tumors. FOXA1 was markedly upregulated in polarized adenocarcinoma cells, where it colocalized with CDH1 (Fig. 4B-D). Conversely, VIM and CD31 were predominantly expressed in non-tumor areas, as confirmed by statistical analysis, suggesting a predominance of the epithelial phenotype in early-stage lung adenocarcinoma cells (Fig. 4B). In human lung adenocarcinoma tissue sections, more than 90% of polarized cells co-expressed FOXA1 and CDH1, a substantial increase compared to the approximately 65% double-positive cells observed in adjacent non-tumorigenic regions, which most of them were AT2 cells (Fig. 4C). This result aligns with similar findings in the mouse model of lung adenocarcinoma induced by *Kras*^G12D^ mutation (Fig. 2D).

In human normal lung tissue, FOXA1 expression was strongly correlated with *CDH1* mRNA expression levels (R = 0.52, p < 0.01), suggesting that FOXA1, as a transcription factor, may play a role in regulating the epithelial state of normal lung AT2 cells (Extended Data Fig. 7A). Gene expression analysis of non-metastatic human NSCLC revealed a significant positive correlation between FOXA1 and CDH1 levels (R = 0.44, p = 0.03; Extended Data Fig. 7B). In contrast, this correlation was not observed in metastatic NSCLC tumors[77], implying that FOXA1 controls CDH1 levels in well-differentiated lung adenocarcinoma cells but not in more aggressive tumors.

Profiling A549 adenocarcinoma cells along the EMT spectrum with ATAC-seq and H3K27ac and FOXA1 CUT&Tag revealed that *CDH1* mRNA expression peaked in the epithelial state. This induction coincided with FOXA1 binding to a H3K27ac-enriched enhancer within the *CDH1* locus, implicating FOXA1 in directly activating *CDH1* transcription in the epithelial state (Extended Data Fig. 7C).

MIHC of human LUAD sections revealed discrete polarized tumor structures (Fig. 4E). Analysis of serial H&E and mIHC sections showed that these foci were enriched for FOXA1 and CDH1 relative to the surrounding stroma. A focal reduction of FOXA1 staining was detectable within one region of the polarized cluster. FOXA1-positive cell collectives exhibited strong CDH1 expression and asymmetric nuclear positioning, hallmarks of epithelial cell polarity, whereas areas with reduced FOXA1 lost LN and CDH1-defined apical–basal organization (Fig. 4E). FOXA1-high epithelial-like LUAD cells were surrounded by VIM-positive stromal cells and α-SMA/MMP2-positive myofibroblasts, whereas LUAD cells with reduced FOXA1 expression exhibited increased MMP2 expression. These polarized, FOXA1-enriched cells also stained positive for KI67, indicating active proliferation. Collectively, the findings suggest that elevated FOXA1 is required to sustain polarized architecture in LUAD.

In advanced human LUAD specimens, FOXA1 expression was significantly diminished in solid tumor regions characterized by reduced cellular differentiation, relative to adjacent polarized areas (Extended Data Fig. 7D), reinforcing the association between elevated FOXA1 expression and epithelial cell polarity in LUAD cells.

### FOXA1-KD reduces intercellular adhesion, proliferation, and tumorigenesis and promotes oxidative phosphorylation in A549 adenocarcinoma cells

To further investigate whether FOXA1 directly regulates cell polarity, proliferation, and tumorigenesis in lung adenocarcinoma cells, CRISPR technology was initially used to generate FOXA1-KD A549 adenocarcinoma cell lines; however, a complete ablation of FOXA1 expression was not achieved (data not shown). As an alternative, a lentiviral-mediated KD approach was adopted. A Tet-on inducible lentiviral system was used to deliver shRNA targeting FOXA1, as previously described, to enable precise assessment of cellular changes before and after FOXA1 suppression[78]. Treatment with doxycycline for 11 days significantly reduced FOXA1 protein levels, as verified by western blot analysis in shFOXA1 #1 and #2 A549 clones using two identical shRNA constructs (Fig. 4F).

FOXA1-KD markedly increased the EMT score in clone shFOXA1#2, indicating that depletion of FOXA1 drives A549 adenocarcinoma cells toward a mesenchymal phenotype (Fig. 4G). Consistent with this finding, gene set enrichment analysis (GSEA) in the FOXA1-deficient H2228 human lung adenocarcinoma cell line demonstrated significant enrichment of the EMT Hallmark gene set in shFOXA1 clones (Supplementary Table 2). The same trend was observed in the murine lung adenocarcinoma cell line LA795, where all three shFOXA1 clones showed significant enrichment of the EMT Hallmark gene set (Supplementary Table 2).

To assess the impact of FOXA1-KD on lung adenocarcinoma cell polarity, immunofluorescence co-staining of actin microfilaments and CDH1 was performed in A549 adenocarcinoma cells treated with B1.14 to enhance their epithelial-like characteristics (Extended Data Fig. 7E). In shNC A549 control cells, actin microfilaments co-localized robustly with CDH1. In contrast, the colocalization was disrupted in FOXA1-KD cells. In FOXA1-KD A549 cells, CDH1 adopted a punctate pattern on the cell surface, indicative of compromised cell polarity[79].

The expression of the cell cycle arrest-related protein P21 was upregulated in FOXA1-KD cells[80], further supporting the role of FOXA1 in driving the cell cycle (Fig. 4H). Furthermore, mTOR expression level was reduced in FOXA1-KD cells. Some studies have demonstrated that mTOR regulates cell proliferation, suggesting that FOXA1 promotes cell cycle activation in lung adenocarcinoma cells[81, 82].

To validate these findings, the cell cycle profiles of control and FOXA1-KD A549 adenocarcinoma cells were analyzed using propidium iodide (PI) staining. PI staining results confirmed that FOXA1 promotes cell proliferation by driving more cells into the S phase while decreasing the proportion of cells in the G1 phase (Extended Data Fig. 7F,G). Previous studies have shown that reduced cell proliferation can increase G0 phase occurrence or enhance cellular senescence (reviewed in[83]). To test whether FOXA1 regulates the cell cycle by inhibiting the G0 phase and cellular senescence, a p27k-mVenus vector[84] was used to transfect shNC and shFOXA1 cells. These cells were then treated with B1.14, DMSO, or TGF-β for 72 hours, and P27 protein expression was assessed via mVenus expression using fluorescent-activated cell sorting (FACS) analysis. The results revealed that cells treated with TGF-β accumulated in the G0 phase compared to B1.14-treated cells (Extended Data Fig. 7H), consistent with the lower tumor size observed following tail vein injection of cells treated with TGF-β (Fig. 1E-G).

RNA-seq profiling of shNC cells and two shFOXA1-KD clones revealed a widespread down-regulation of proliferation-related genes in the shFOXA1 samples (Fig. 4I). Additionally, cellular senescence was significantly elevated in the shFOXA1 A549 adenocarcinoma cells, as evaluated by β-galactosidase staining (Fig. 4J).

To further investigate whether A549 adenocarcinoma cells with reduced FOXA1 expression could exhibit the same phenotype in vivo, 1 million shNC, shFOXA1#1, and shFOXA1#2 A549 adenocarcinoma cells were injected into the tail vein of immunodeficient NCG mice. Lung tissue was evaluated after five weeks (Fig. 4K). Consistent with in vitro findings, A549 adenocarcinoma cells with low FOXA1 expression displayed reduced tumor forming potential (Fig. 4L,M).

In *Kras*^G12D^ driven murine NSCLC, FOXA1-KD tumors displayed up-regulated metabolic pathways and heightened oxidative phosphorylation (Fig. 3J). These results were confirmed by assessing the glycolytic flux and mitochondrial stress in shNC and shFOXA1 clones. FOXA1 silencing increased glycolysis and enhanced mitochondrial stress, mirroring the in vivo observations (Extended Data Fig. 7I,J).

### Reducing FOXA1 expression in lung adenocarcinoma cells restores lysosome-like organelles

As previously shown, *Kras*^G12D^ lung adenocarcinoma cells with FOXA1-KD accumulate lysosome-like organelles, and spatial transcriptomics revealed concurrent up-regulation of lysosome-associated genes and down-regulation of genes involved in lysosomal membrane fusion in these FOXA1-KD malignant cells (Extended Data Fig. 3A, Fig. 3K). Apical-basal polarized cells often exhibit an elongated cell morphology[85, 86]. The structural characteristics of lung adenocarcinoma cell apical-basal polarity were examined under FOXA1 regulation. Specifically, the abnormal proliferation of AT2 cells and the differences between normal AT2 cells and polarized lung adenocarcinoma cells were investigated.

The morphology of normal AT2 cells, proliferating AT2 cells, abnormally proliferating AT2 cells, and polarized lung adenocarcinoma cells in *Kras*^G12D^ mice before and after Sftpc-Cre induction was analyzed by TEM (Fig. 5A). One of the most critical features of normal AT2 cells is an abundance of lamellar bodies and lysosome organelles that generate and secrete surfactant proteins into the alveoli[87, 88] (Fig. 5A). In polarized regions of lung adenocarcinoma, cells displayed a distinct basement membrane, with nuclei positioned adjacent to it, and remained closely apposed through junctional complexes. Interestingly, the most striking observation was a dramatic decrease in lysosome-like organelles in polarized lung adenocarcinoma cells compared to unpolarized carcinoma cells and normal AT2 cells (Fig. 5A).

**Fig. 5.**
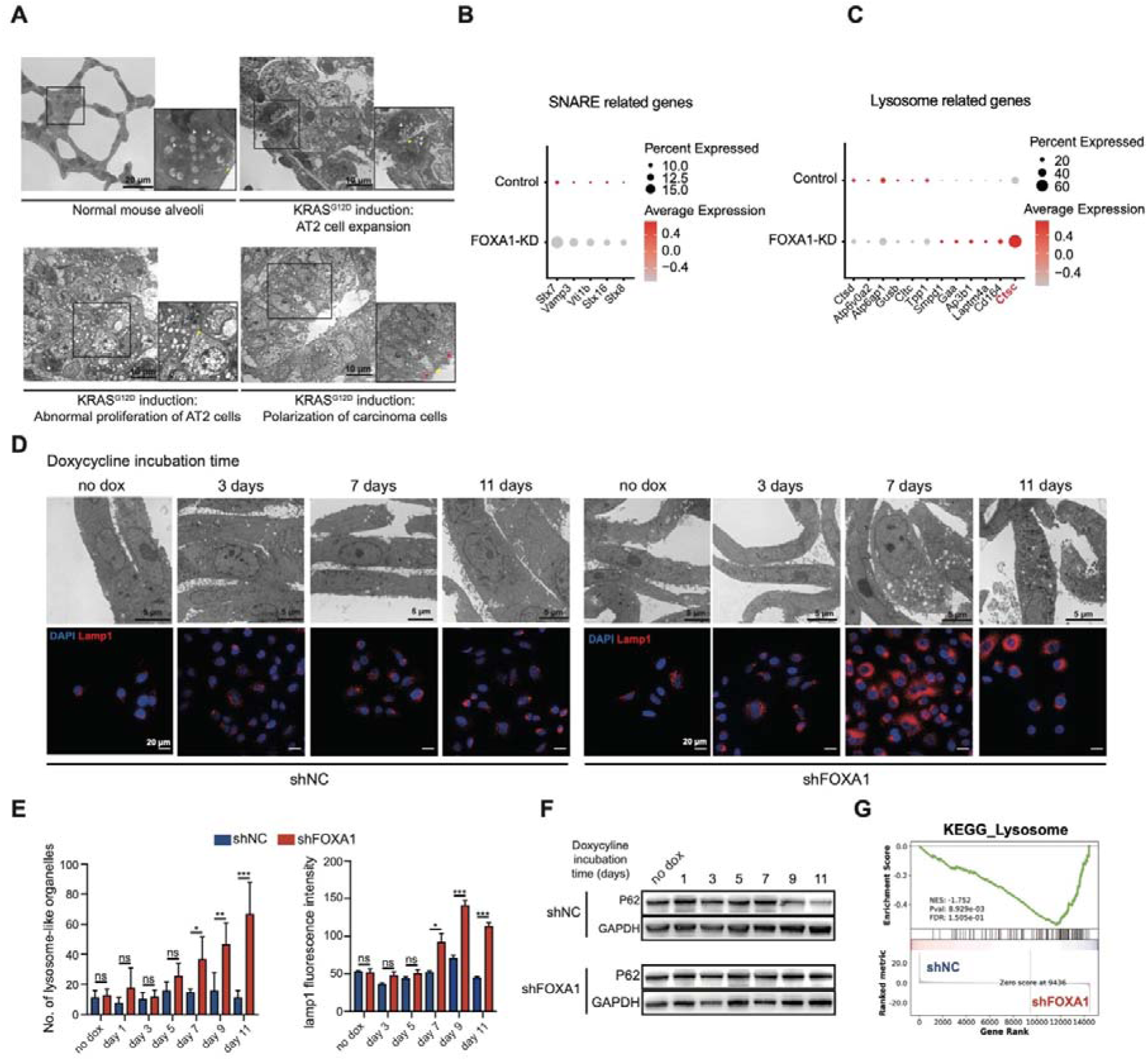
Lysosomes are restored in lung adenocarcinoma cells with reduced FOXA1 expression. **A**, Electron microscopy of a normal mouse lung and *Kras*^G12D^ driven (90 days) lung adenocarcinoma tumors. **B**, Expression of SNARE-associated genes in control and FOXA1-KD malignant cells from Visium HD spatial analysis. **C**, Expression of Lysosome-associated genes in control and FOXA1-KD malignant cells from Visium HD spatial analysis. **D**, Electron microscopy and LAMP-1 immunofluorescence images of A549 lung adenocarcinoma cells expressing shNC or shFOXA1 at successive days following doxycycline induction. **E**, Bar plots demonstrated an increase in both the number of lysosome-like organelles and LAMP-1 fluorescence following FOXA1-KD. Statistical analysis was conducted using t-test. **F**, Western blot result of P62 and GAPDH on shNC and shFOXA1 A549 adenocarcinoma cells. **G**, KEGG analysis of lysosome-related genes on RNA-seq result of shNC and shFOXA1 A549 adenocarcinoma cells.

Pathway analysis of malignant cells in FOXA1-KD *Kras*^G12D^ versus control mice revealed a downregulation of SNARE gene expression (Fig. 3K). The expression levels of STX7, STX8, STX16, VAMP3, and VTI1B were significantly downregulated in FOXA1-KD *Kras*^G12D^ malignant cells compared to control (Fig. 5B). STX7, STX8, and STX16, all members of the SNARE protein family, play critical roles in lysosome membrane fusion processes with phagosomes. The downregulation of SNARE genes in FOXA1-KD tumors may account for the accumulation of lysosomes and vacuolar structures observed by electron microscopy in LUAD cells.

Additionally, the expression of several lysosome-associated genes was found to be increased in FOXA1-KD *Kras*^G12D^ mice malignant cells, with a marked upregulation of cathepsin C (CTSC) (Fig. 5C). CTSC has been shown to promote breast cancer metastasis to the lung by modulating neutrophil function and enhancing the formation of neutrophil extracellular traps (NETs)[89]. Furthermore, CTSC plays a pivotal role in the progression of NSCLC by promoting cell motility, and invasiveness. Overexpression of CTSC in NSCLC cells enhances their malignant potential both in vitro and in vivo, whereas silencing CTSC expression reverses this phenotype. Mechanistically, CTSC promotes EMT through activation of the Yes-associated protein (YAP) signaling pathway[90]. These studies were consistent with our findings in *Kras*^G12D^ FOXA1-KD malignant cells, which exhibit enhanced malignancy and an elevated EMT phenotype.

Since polarized lung adenocarcinoma cells exhibit high FOXA1 expression, FOXA1 may influence lysosome biogenesis in these cells. To investigate this, a doxycycline-inducible TET-on system was utilized to express FOXA1-targeting shRNA, as previously described. FOXA1 protein levels were significantly reduced after 3–5 days of doxycycline treatment (data not shown). Before doxycycline induction, there was no significant difference in the number of lysosome-like organelles between shNC and shFOXA1 A549 adenocarcinoma cells (Fig. 5D,E). Immunofluorescence staining for lysosomal-associated membrane protein 1 (LAMP1), a lysosomal marker[91, 92], also showed no differences before doxycycline treatment (Fig. 5D,E). However, following doxycycline treatment, a gradual increase in the number of lysosomes and LAMP1 was observed by immunofluorescence over time in the FOXA1-deficient cells (Fig. 5D,E). We also observed the same increase in the human RERFLCMS lung adenocarcinoma cells and mouse LA795 lung adenocarcinoma cells (Extended Data Fig. 8A-D).

Doxycycline has been reported to act as a cellular stressor, influencing autophagy flux in carcinoma cells[93]. It functions as an activator of autophagy, promoting autophagy flux and lysosome regeneration. To evaluate autophagy flux, P62 protein levels were examined by western blotting (Fig. 5F). In A549 adenocarcinoma cells with normal FOXA1 expression, P62 protein levels decreased gradually with increasing incubation time under doxycycline treatment, reflecting active autophagy flux. However, in FOXA1-KD A549 adenocarcinoma cells, P62 protein levels did not decrease after doxycycline induction, indicating a disruption of autophagy flux in shFOXA1 cells. This finding paralleled the results in *Kras*^G12D^ murine lung tumors, where the control tumors displayed elevated autophagic activity relative to the FOXA1-KD *Kras*^G12D^ tumors (Fig. 3K).

The lysosome biogenesis pathway was further analyzed using GSEA, revealing that shFOXA1 A549 adenocarcinoma cells exhibited higher lysosome activity than the control cells (Fig. 5G). Additionally, several SNARE proteins were downregulated in shFOXA1 cells (Extended Data Fig. 8E). These data suggest that shFOXA1 A549 adenocarcinoma cells exhibit enhanced lysosomal biogenesis leading to an abnormal lysosomal accumulation.

Previous studies have demonstrated that lamellar bodies, lysosome-related organelles in alveolar type II cells, chiefly mediate surfactant protein secretion and preserve alveolar architecture[94, 95]. FOXA1 can bind to the promoter region of surfactant protein B to upregulate its expression, while such binding could be inhibited through Smad3 by TGF-β signaling pathway induction[96]. During lung development, FOXA1 regulates AT2 cell maturation[97].

Overall, our results demonstrate that proliferating, polarized human adenocarcinoma cells possess few lysosomes, whereas FOXA1-KD lung adenocarcinoma cells harbor large number of lysosomes through the activation of lysosome biogenesis and suppression of lysosome-autophagosome fusion. This pattern was consistent with the increased expression of lysosomal genes, such as CTSC, and the marked reduction in SNARE gene expression observed in *Kras*^G12D^ mouse models following FOXA1-KD.

### FOXA1 binds competitively with TFE3 in lysosome-encoding loci and disrupts lysosome biogenesis in polarized A549 adenocarcinoma cells

FOXA1 negatively regulates lysosome regeneration in polarized lung adenocarcinoma cells. To investigate the mechanism underlying lysosome regeneration defects in these cells, Ingenuity Pathway Analysis (IPA) was performed using the list of upregulated genes in FOXA1-KD A549 adenocarcinoma cells compared to control cells. IPA identified the Coordinated Lysosomal Expression and Regulation (CLEAR) signaling pathway as the top pathway associated with lysosome regeneration following FOXA1-KD (Fig. 6A). The CLEAR signaling pathway is a crucial cellular mechanism that regulates gene expression in lysosomal biogenesis and function. This pathway is primarily controlled by the MiT/TFE family of transcription factors, which includes Microphthalmia-associated Transcription Factor (MITF), Transcription Factor EB (TFEB), and TFE3[98, 99].

**Fig. 6.**
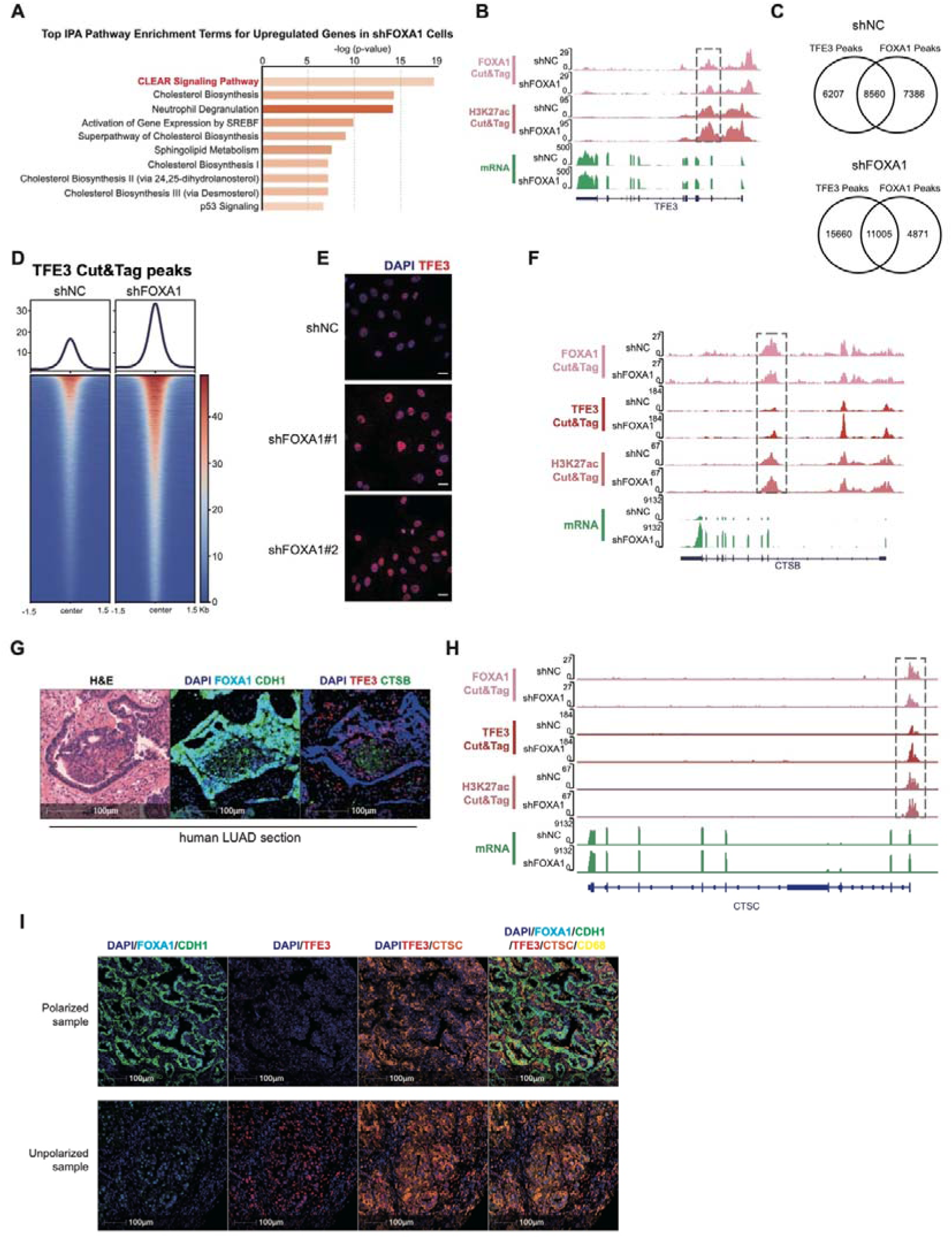
FOXA1 competes with TFE3 at lysosomal gene loci, leading to a reduction in lysosomal organelles in human adenocarcinoma cells and LUAD tissues. **A**, IPA analysis of upregulated pathways in shFOXA1 cells compared to shNC A549 adenocarcinoma cells. **B**, IGV of FOXA1 and H3K27ac Cut&Tag peak as well as TFE3 mRNA expression on the TFE3 locus in shNC and shFOXA1 A549 adenocarcinoma cells. **C**, Cut&Tag peak overlaps of FOXA1 and TFE3 in shNC and shFOXA1 A549 adenocarcinoma cells. **D**, Heatmap of TFE3 Cut&Tag peaks in shNC and shFOXA1 A549 adenocarcinoma cells. **E**, Immunofluorescence staining of TFE3 on shNC and shFOXA1#1 and shFOXA1#2 A549 adenocarcinoma cells. **F**, IGV tracks of FOXA1, TFE3, and H3K27ac Cut&Tag peaks, along with CTSB mRNA expression at the CTSB locus, in shNC- and shFOXA1-expressing A549 adenocarcinoma cells. **G**, Serial sections of human LUAD tissue stained with H&E, followed by mIHC panels comprising DAPI/FOXA1/CDH1 and, on adjacent sections, DAPI/TFE3/CTSBs staining. **H**, IGV peaks of FOXA1, TFE3 and H3K27ac Cut&Tag peaks as well as CTSC mRNA expression on CTSC locus in shNC and shFOXA1 A549 adenocarcinoma cells. **I**, Polarized and unpolarized human LUAD tissue staining with mIHC panels comprising DAPI/FOXA1/CDH1/TFE3/CTSC/CD68.

TFE3, a key regulator of the CLEAR signaling pathway, was significantly upregulated in shFOXA1 A549 adenocarcinoma cells. FOXA1 Cut&Tag and H3K27ac Cut&Tag analyses revealed that FOXA1-KD resulted in diminished FOXA1 occupancy and increased H3K27ac enrichment at TFE3 enhancer regions. These results indicated that diminished FOXA1 binding is associated with attenuated FOXA1-dependent regulation of TFE3 in the shFOXA1 A549 adenocarcinoma cells. Furthermore, the increased enhancer activity and upregulated TFE3 transcription following FOXA1-KD suggest that TFE3 expression becomes FOXA1-independent under these conditions (Fig. 6B). TFE3 serves as a transcription factor regulating lysosomal biogenesis, suggesting that TFE3 represents a primary downstream target of FOXA1 in the pathway mediating the reduction of lysosome abundance in polarized lung adenocarcinoma cells.

Cut&Tag assays targeting TFE3 in shNC and shFOXA1 A549 adenocarcinoma cells were performed to address this hypothesis. Compared to shNC cells, shFOXA1 cells exhibited elevated TFE3 binding peaks (Fig. 6C). Additionally, the density of TFE3 peaks was significantly higher in the shFOXA1 cells (Fig. 6D). These findings indicate that FOXA1-KD enhances TFE3 activity and its role in regulating the CLEAR signaling pathway, which contributes to lysosome regeneration in A549 adenocarcinoma cells.

To further investigate the role of TFE3 in DNA binding and transcriptional regulation after FOXA1-KD, immunolabeling of TFE3 was performed in shNC and shFOXA1 A549 adenocarcinoma cell lines. Higher TFE3 protein levels were found in the nuclei of shFOXA1 cells compared to the shNC cells (Fig. 6E). CTSB, a known target of TFE3, is a lysosomal cysteine protease[100, 101]. CTSB expression was significantly increased following FOXA1-KD. The enhancer region of CTSB exhibited a higher TFE3 binding peak in the shFOXA1 group than the shNC cells, while FOXA1 binding was reduced in the same enhancer region defined by H3K27ac peaks with significantly higher mRNA expression in shFOXA1 condition (Fig. 6F). These findings suggest that in tumors with polarized adenocarcinoma cells that overexpress FOXA1, CTSB activity may be impaired, leading to a diminished ability to generate functional lysosomes.

MIHC co-staining of FOXA1/CDH1 and TFE3/CTSB was conducted on two adjacent sections of human lung adenocarcinoma (LUAD) tissue samples (Fig. 6G). As expected, in the most polarized regions where FOXA1 and CDH1 expressions were elevated, TFE3 had a reduced expression in the nucleus, and CTSB was nearly absent (Fig. 6G). In contrast, in unpolarized regions with low FOXA1 expression, higher TFE3 levels were observed in the nucleus, along with increased CTSB expression (Fig. 6G). Another lysosome-associated gene, CTSC, was initially found to be highly expressed in *Kras*^G12D^ FOXA1-KD malignant cells (Fig. 6C) and was also regulated by TFE3 at the regulatory region in human A549 adenocarcinoma cells together with H3K27ac binding (Fig. 6H). Elevated CTSC protein expression was detected predominantly in the unpolarized regions, as revealed by mIHC co-staining for FOXA1, CDH1, TFE3, CTSC, and CD68 on human LUAD samples encompassing both polarized and unpolarized regions (Fig. 6I). Notably, in the unpolarized regions, where TFE3 exhibited strong nuclear localization, FOXA1 expression was markedly reduced.

Collectively, we uncovered a marked reduction in lysosome abundance, a previously unrecognized phenotype in polarized lung adenocarcinoma cells. Mechanistically, FOXA1 competes with TFE3 for binding to the regulatory elements of lysosome-biogenesis genes, especially CTSB and CTSC, thereby curtailing their transcription. Concordantly, lysosome numbers decline as NSCLC cells proliferate and polarize in concert with FOXA1 upregulation.

## Discussion

In this study, we reveal a pivotal role for FOXA1 in preserving epithelial cell polarity and suppressing lysosome biogenesis in NSCLC. Our results demonstrate that FOXA1 is highly expressed in polarized, well-differentiated lung adenocarcinoma cells in human tumors and in the *Kras*^G12D^ mouse model. FOXA1 is essential for maintaining epithelial identity, facilitating proliferation, and supporting robust tumorigenesis. Conversely, loss or reduction of FOXA1 expression leads to the acquisition of mesenchymal features, increased lysosome biogenesis, impaired proliferation, and significant enhance TGF-β signaling on cells throughout the microenvironment.

Our data reinforce the concept that FOXA1 serves as a guardian of the epithelial state in lung adenocarcinoma. Through comprehensive epigenomic profiling (ATAC-seq, CUT&Tag) and functional assays, we show that FOXA1 occupies chromatin regions associated with epithelial gene expression, notably CDH1, and its presence correlates with strong epithelial signatures. Importantly, FOXA1-KD, both in vitro and in vivo, induces EMT-like changes, characterized by decreased CDH1 and enhanced TGF-β signaling activity. These findings are consistent with previous reports implicating TGF-β signaling as a master regulator of EMT and tumor progression in NSCLC, and position FOXA1 as a critical counterbalance to this pathway[4].

A novel aspect of our work is the elucidation of the function of FOXA1 in regulating lysosome biogenesis. Our results reveal that polarized, FOXA1^high^ lung adenocarcinoma cells exhibit a striking reduction in lysosomal abundance. Mechanistically, we uncover that FOXA1 represses lysosome biogenesis by competing with TFE3 for binding to regulatory elements of lysosome-associated genes, including CTSC and CTSB. FOXA1 loss leads to increased TFE3 activity and transcriptional upregulation of lysosomal genes, resulting in enhanced lysosome formation and disrupted autophagic flux. In addition, FOXA1-KD lung malignant cells reduce the expression of SNARE proteins, which also leads to the accumulation of lysosomes in those cells. STX7, a lysosome-associated Qa-SNARE, is essential for the direct fusion of late endosomes with lysosomes, a key step in lysosome formation[102, 103]. Similarly, STX8 is localized to lysosomes and endosomal compartments, functioning as a Qc-SNARE required for various membrane fusion events involved in lysosome biogenesis. STX8 forms complexes with STX7, VTI1B, and other SNAREs to mediate late endosome–lysosome fusion[103]. Loss of STX8 results in defects in lysosome-associated compartments and impaired cellular function, underscoring its essential role in lysosome formation[104]. STX16 is involved in autophagy-related membrane fusion, facilitating the trafficking of ATG9a vesicles to autophagosomes and promoting autolysosome formation[105]. VAMP3, a v-SNARE protein predominantly localized to recycling endosomes and the plasma membrane, regulates vesicular trafficking and membrane fusion events across various cell types[106]. This axis links polarity, cell state, and intracellular organelle dynamics in lung cancer cells, suggesting that FOXA1-mediated suppression of lysosomal biogenesis is integral to maintaining the epithelial, proliferative phenotype.

Lysosomes have recently been identified as important mediators of the proto oncogene tyrosine protein kinase SRC delivery to the cell surface through fusion with autophagosomes to form autolysosomes, followed by exocytotic release of SRC of carcinoma cells [107]. This newly described mechanism reveals a previously unrecognized role of lysosomes in cancer invasion and metastasis and underscores their importance in tumor progression. In this study, we demonstrated that advanced-stage lung adenocarcinoma with reduced polarization capacity exhibits loss of FOXA1 binding and enhanced TFE3-mediated regulation, which increases lysosome abundance in lung carcinoma cells.

From a translational perspective, our findings have several important implications. First, we identify FOXA1 as a potential biomarker for tumor differentiation status and prognosis in lung adenocarcinoma, as its expression is markedly reduced in poorly differentiated, high-grade tumors. Second, the FOXA1–TFE3–lysosome axis may offer therapeutic opportunities; targeting lysosome biogenesis or restoring FOXA1 function could make chemotherapy more effective and limit tumor aggressiveness. Third, modulation of FOXA1 levels may influence the TGF-β signaling activity in cells across the TME.

There are some limitations to our study. Although we establish correlative and functional links between FOXA1, polarity, and lysosome biogenesis, further work is necessary to decipher the precise molecular mechanisms of FOXA1–TFE3 competition and to validate these pathways in larger clinical cohorts. Additionally, our models primarily focus on early-stage, well-differentiated lung adenocarcinoma; the role of FOXA1 in metastatic or therapy-resistant disease remains to be fully explored.

### Conclusions

In conclusion, our work reveals a central role for FOXA1 in maintaining epithelial polarity and regulating lysosome biogenesis in lung adenocarcinoma. These findings enhance our understanding of the molecular determinants of tumor cell state and plasticity, and they open new avenues for therapeutic intervention in NSCLC.

## Declarations

### Ethics, Consent to Participate, and Consent to Publish declarations

Ethics, data availability, all animal experiments were conducted in accordance with institutional guidelines and approved by the Guangzhou Medical University IACUC (protocol S2022-508), and patient samples were obtained with informed consent and institutional IRB approval. All participants provided written informed consent prior to enrollment. The study was reviewed and approved by the Institutional Ethics Committee of the First Affiliated Hospital of Guangzhou Medical University (Access Number: 2022-71) and was conducted in accordance with the Declaration of Helsinki.

## Data availability

scRNA-seq, Visium HD data, RNA-seq data, ATAC-seq and Cut&Tag sequencing data are available from the National Genomics Data Center (https://ngdc.cncb.ac.cn/) under accession number subCRA050457. The remaining data are available within the article and extended data or from the corresponding authors upon request.

## Acknowledgments

We thank GIBH animal facility for animal care and animal experience. Flow cytometry analysis was performed at Flow Cytometry Facility at Guangzhou laboratory. EM analysis was performed at the EM Facility at Pearl River Fisheries Research Institute (PRFRI) with the help of Ms. Chang Ouqin and Ms. Chen Jiayue. scRNA-seq and HD visium analysis were performed by the NovelBio team. We thank Yan Xin, Xie Mengze, Chen Chunke and Yuan Tingjie for their technical support.

## Author contributions

J.P.T, J.H. and X.W. conceptualized the study. X.W., B.Z. C.S. and J.P.T. designed the experiments and analyzed the results. B.Z., M.H., W.H., B.Z., X.Z., X.R., G.Z., and Y.Z. performed the experiments. Y.Z., H.L., W.L., M.D.T., T.Z.T., Y.T.Z, helped in conceptualization and experimental planning. X.W. and J.P.T. wrote the paper. All authors edited and reviewed the final manuscript.

---

**Extended data Fig. 1.**
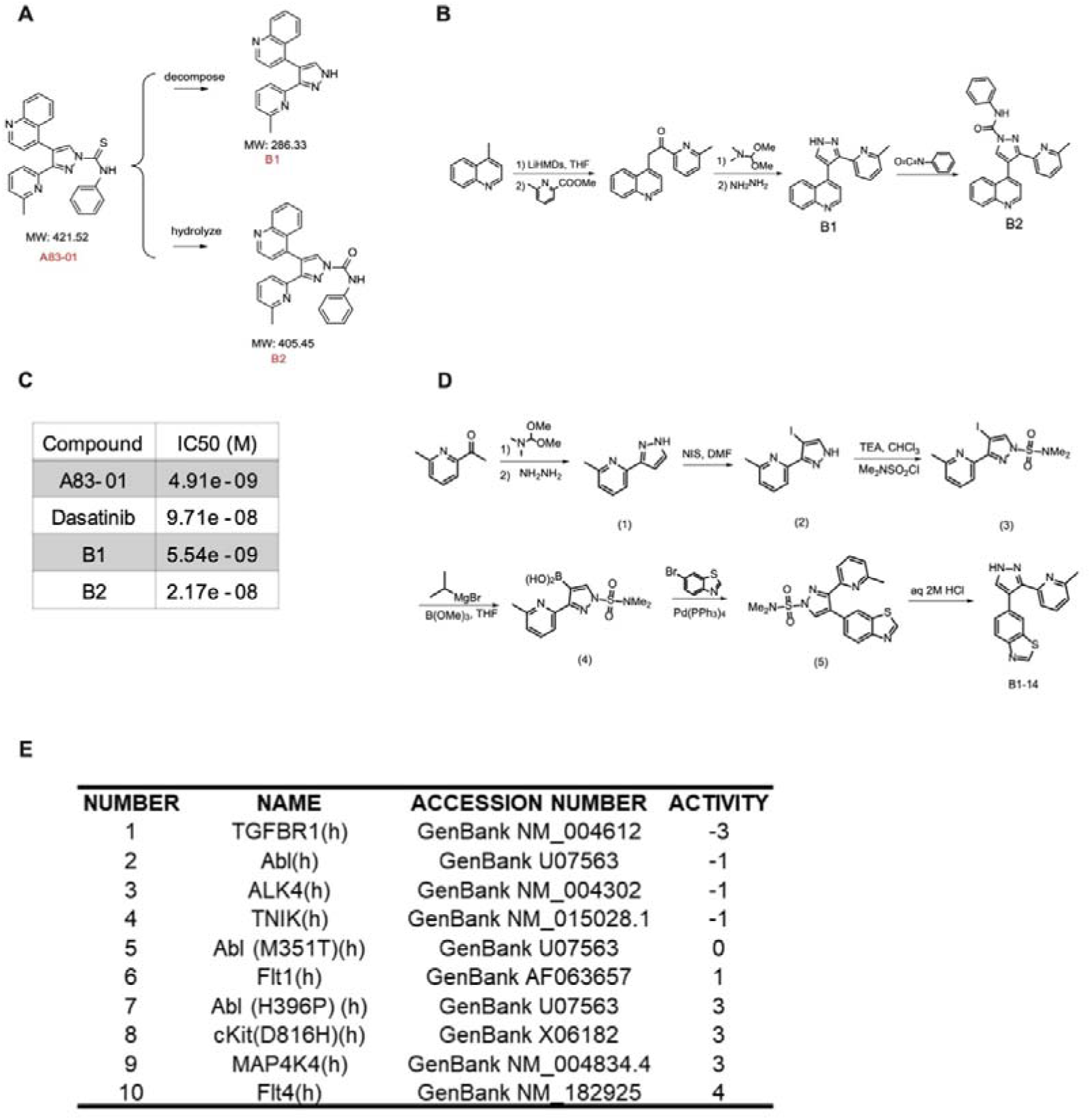
Synthesis of the TGF-β Receptor Inhibitor B1.14 and evaluation of its Kinome Activity. **A**, Pathway of A83-01 breakdown. **B**, Organic synthesis of B1 and B2. **C**, Screening of B1 and B2 and A83-01 against ALK5. **D**, Organic synthesis of B1-14 analog. **E**, Kinase selectivity result shows top 10 reactivity enzymes that reacted with B1.14.

**Extended data Fig. 2.**
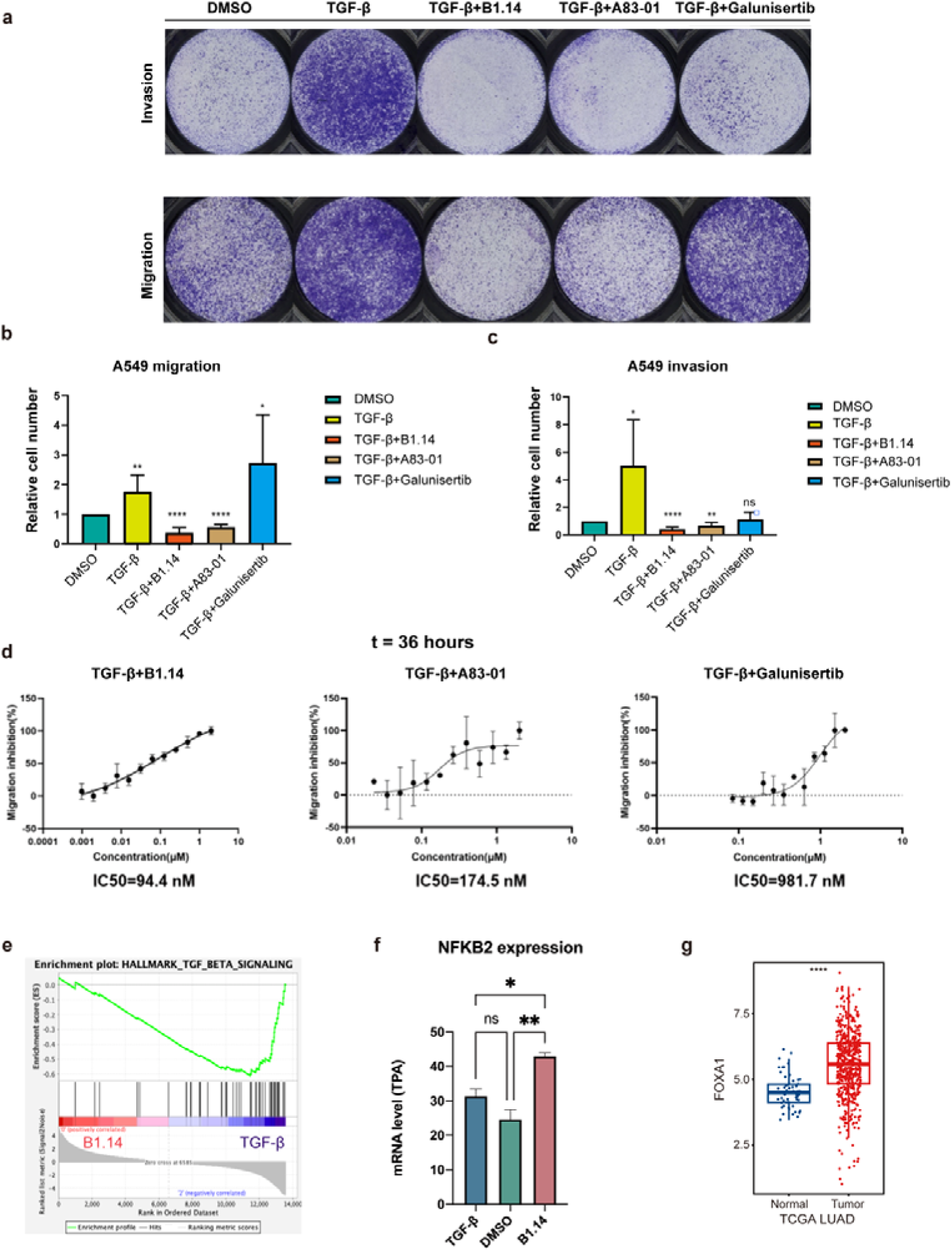
B1.14 inhibits invasion and migration of A549 lung adenocarcinoma cells. **A**, Invasion and migration analysis of A549 cells under treated with DMSO, TGF-β and 3 other TGF-β receptor inhibitors B1.14, A83-01 and Galunisertib. **B&C**, Statistical analysis of A549 cell number passed through chamber membrane in migration (b) and invasion (c) test. Statical analysis was performed using one-way ANOVA. **D**, IC50 of B1.14, A83-01 and Galunisertib calculated by wound healing analysis. **E**, GSEA analysis of Hallmark TGF-β signaling in B1.14 and TGF-β treated A549 cells. **F**, mRNA level of NFKB2 in A549 cells treated with DMSO, B1.14 and TGF-β. Statistical analysis uses one-way ANOVA. **G**, TCGA LUAD data of FOXA1 expression in tumor and normal samples. Statistical analysis is performed using Wilcox-test.

**Extended data Fig. 3.**
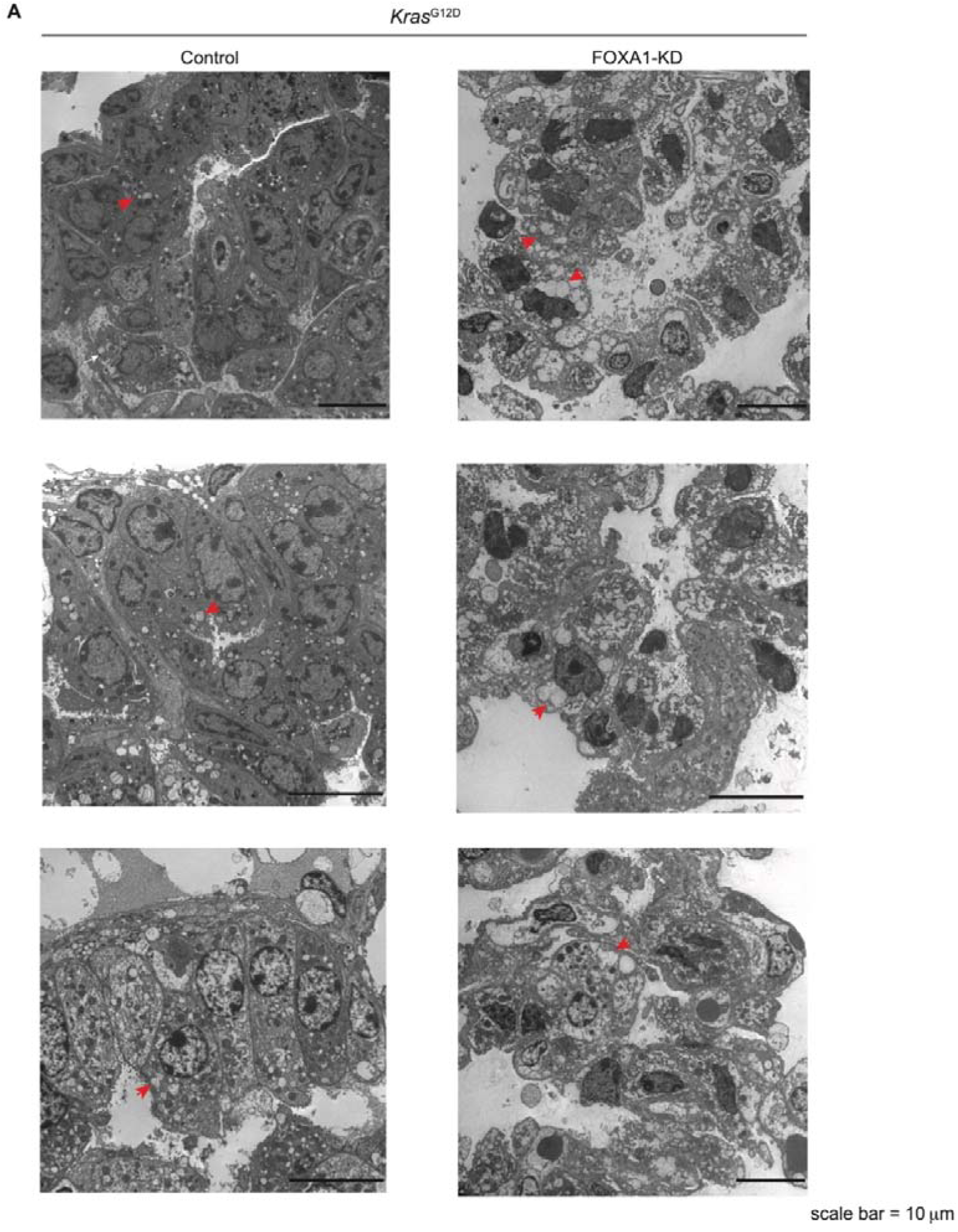
Storage of lysosome-like organelles in FOXA1-KD mouse lung adenocarcinoma cells. **A**, Representative electron microscope images of *Kras*^G12D^ driven lung adenocarcinomas collected 30 days after Cre-mediated activation in control (FOXA1^+/+^) and FOXA1-knockdown (FOXA1^fl/+^) mice. Red arrows indicate lysosome-like organelles (derived from lamellar bodies). Scale bar 10 μm.

**Extended data Fig. 4.**
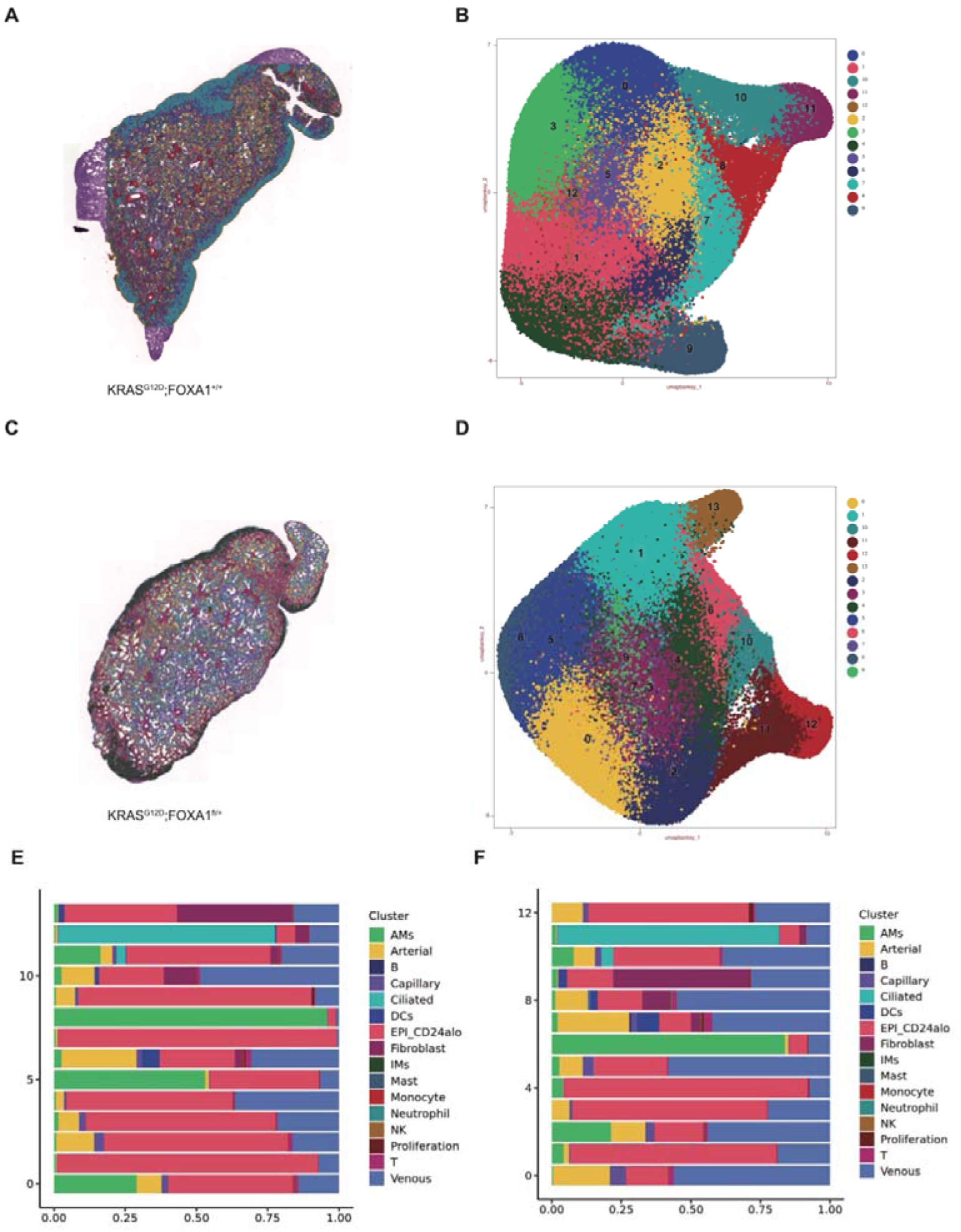
Cellular annotation of spatial analysis. **A&B**, The tissue section (a) and UMAP (b) depicting the unbiased clusters of spatial transcriptomics (ST) spots in *Kras*^G12D^; FOXA1^+/+^ (Control) group. **C&D**, The tissue section (c) and UMAP (d) depicting the unbiased clusters of spatial transcriptomics (ST) spots in *Kras*^G12D^; FOXA1^fl/+^ (FOXA1-KD) group. **E**, Bar plots showing the compositions of cell types in each cluster in Control group. **F**, Bar plots showing the compositions of cell types in each cluster in FOXA1-KD group.

**Extended data Fig. 5.**
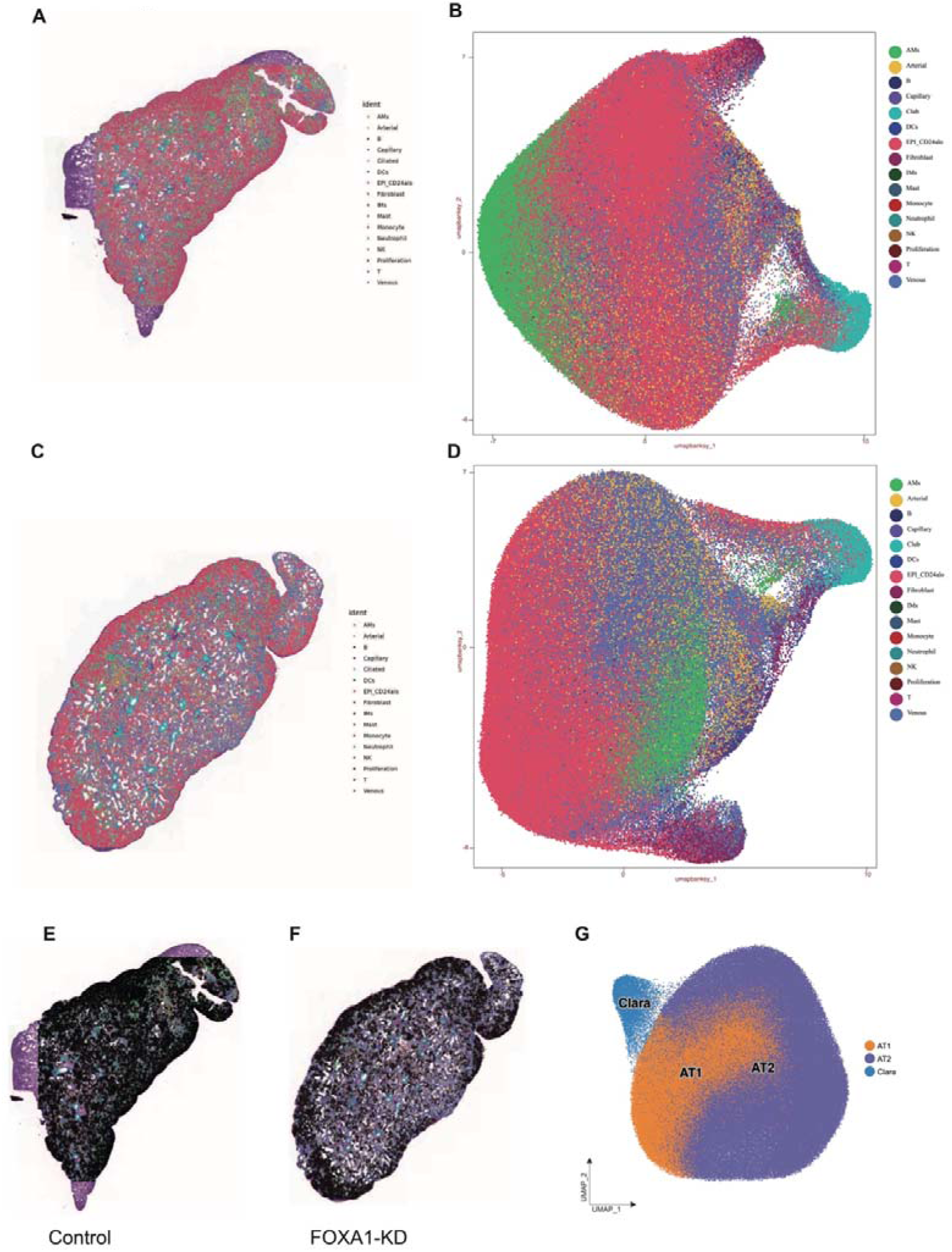
Cellular distribution of spatial analysis. **A&B**, The tissue section and UMAP depicting the annotated cell types of spatial transcriptomics (ST) spots in Control group. **C&D**, The tissue section and UMAP depicting the annotated cell types of spatial transcriptomics (ST) spots in FOXA1-KD group. **E&F**, Distribution of epithelial cells (black) in Control and FOXA1-KD group. **G**, UMAP of AT1, AT2 and Clara cells distribution of Control and FOXA1-KD group.

**Extended data Fig. 6.**
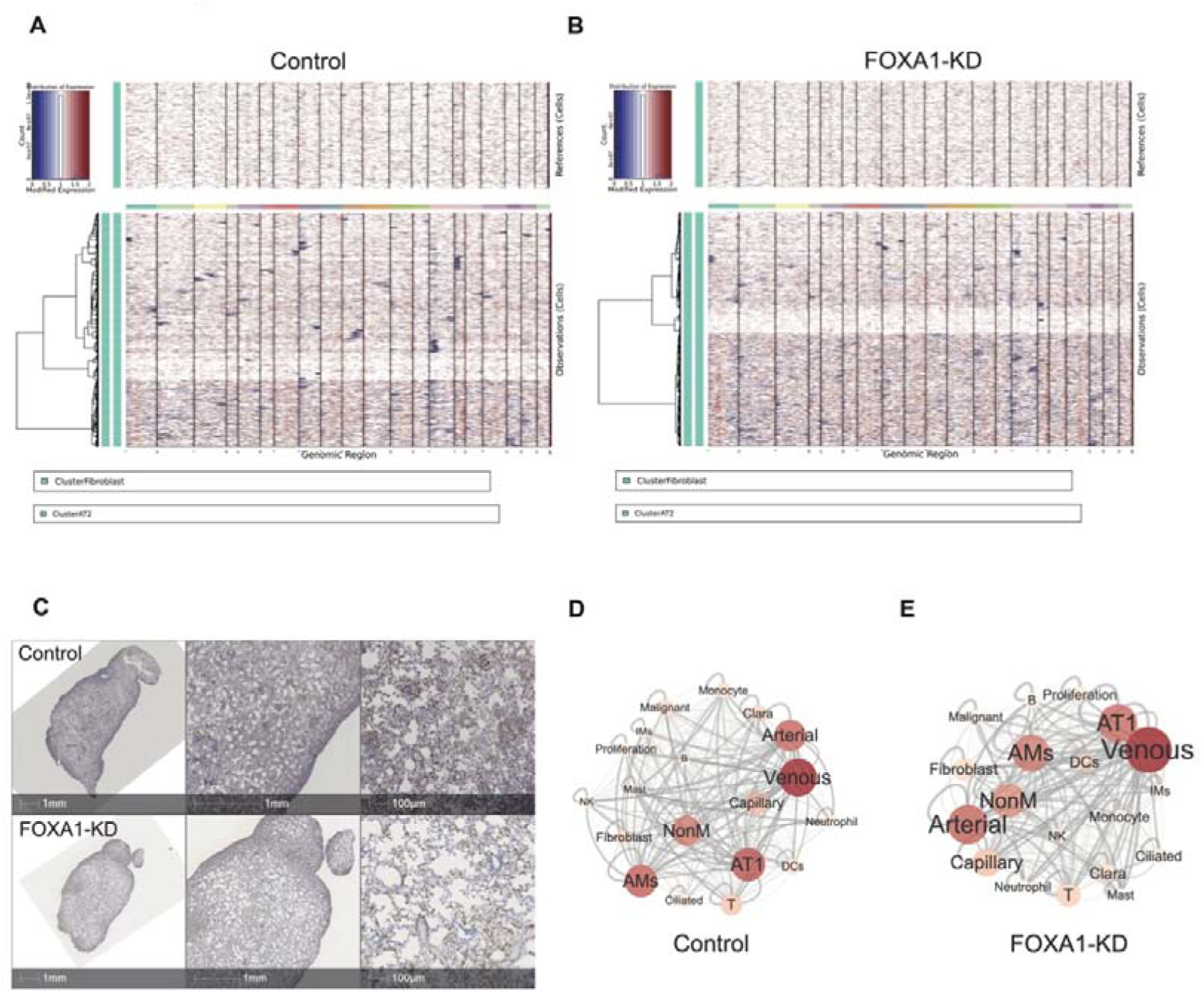
Define malignant cells and analyze the cellular co-occurrence between the control and FOXA1-KD groups. **A&B**, InferCNV calculation of malignant cells in Control (a) and FOXA1-KD (b) group. c, IHC of FOXA1 staining in Control and FOXA1-KD section. **D&E**. Co-occurrence of cell types in Control (d) and FOXA1-KD (e) group.

**Extended data Fig. 7.**
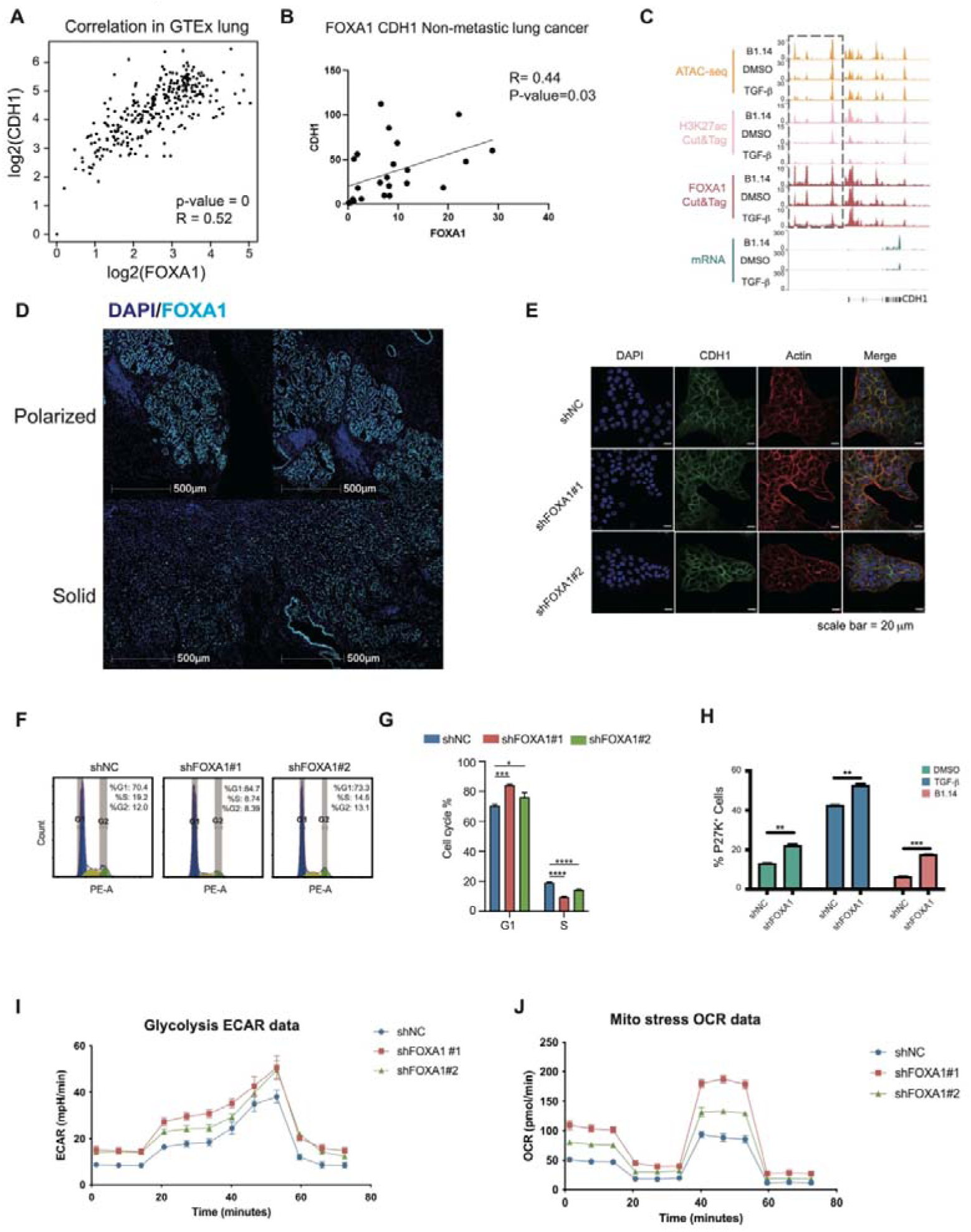
FOXA1-KD lung adenocarcinoma cells exhibited reduced CDH1 expression, prolonged cell cycle duration, an enhanced quiescence-associated profile, and elevated metabolic activity signatures. **A**, Correlation study of CDH1 and FOXA1 expression in GTEx lung dataset. **B**, Correlation study of CDH1 and FOXA1 expression in human non-metastasis NSCLC samples. **C**, IGV tracks of ATAC-seq, H3K27ac Cut&Tag, FOXA1 Cut&Tag, and CDH1 mRNA expression in A549 cells treated with DMSO, TGF-β, or B1.14. **D**, mIHC staining of DAPI and FOXA1 in polarized and solid tumor region of a same late-stage LUAD patient. **E**, Immunofluorescence staining of DAPI, E-cadherin and Actin on shNC, shFOXA1#1 and shFOXA1#2 A549 cells. **F**, FACS analysis of PI staining in shNC, shFOXA1#1 and shFOXA1#2 A549 cells. **G**, Barplot of cell distribution in G1 or S cell cycle in shNC, shFOXA1#1 and shFOXA1#2 A549 cells. Statistical analysis is performed using one-way ANOVA. **H**, Barplot of proportion of P27K+ cells calculated using FACS analysis in shNC and shFOXA1 A549 cells under treated by DMSO, TGF-β, or B1.14. Statistical analysis is performed using one-way ANOVA. **I**, Seahorse glycolysis test of shNC, shFOXA1#1 and shFOXA1#2 A549 cells. **J**, Seahorse mito stess test of shNC, shFOXA1#1 and shFOXA1#2 A549 cells.

**Extended data Fig. 8.**
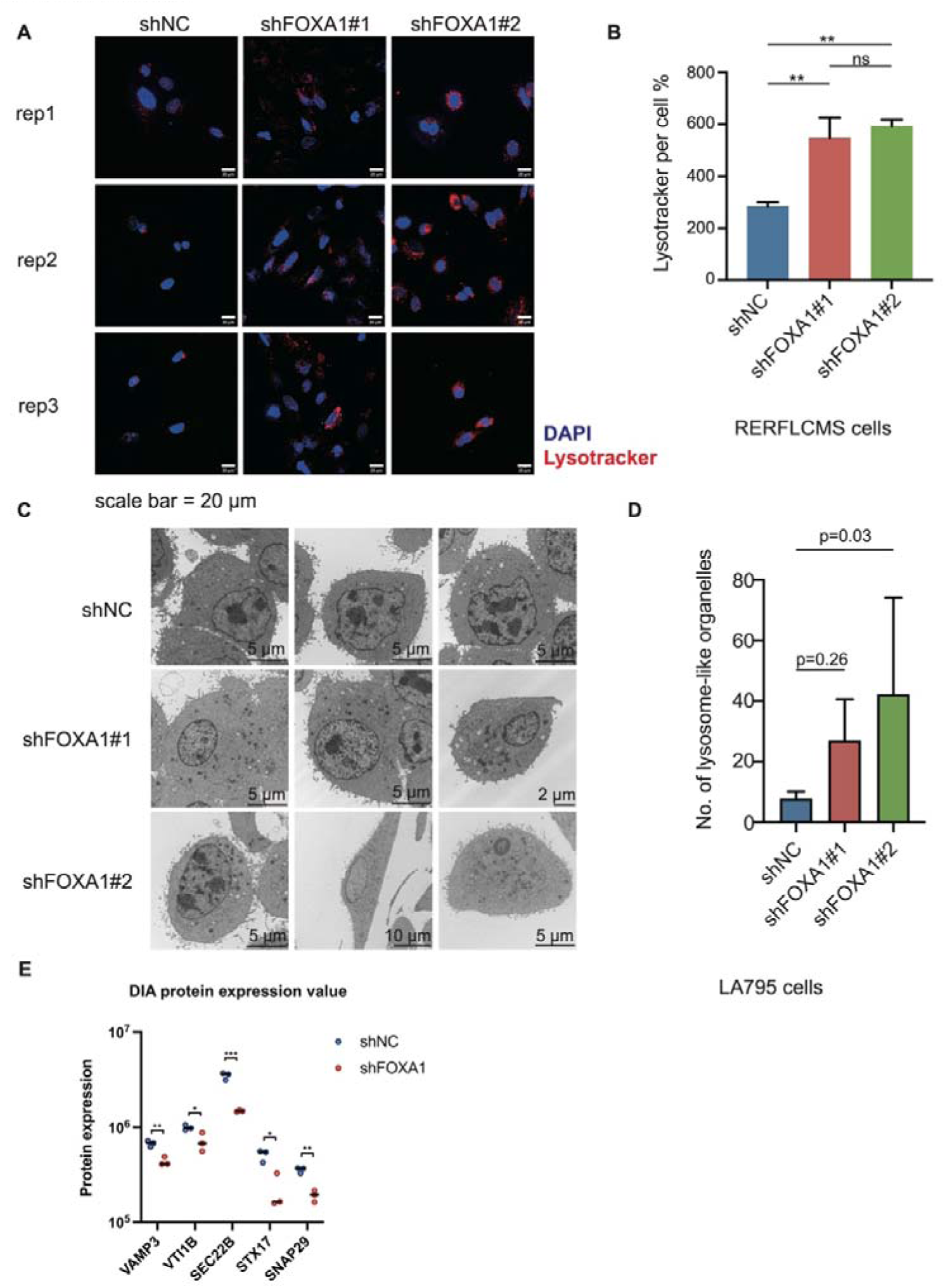
FOXA1 knockdown increases the number of lysosomes in both human and mouse lung adenocarcinoma cell lines. **A**, Lysotracker staining of shNC, shFOXA1#1 and shFOXA1#2 human RERFLCMS cells. **B**, barplot of Lysotracker fluorescence per cell calculated using imageJ software and statistical analysis is performed using one-way ANOVA. **C**, Transmission electron microscope pictures of shNC, shFOXA1#1 and shFOXA1#2 mouse LUAD LA795 cells. **D**, Bar chart illustrating the number of lysosome-like organelles per cell in LA795 mouse LUAD cells expressing shNC, shFOXA1#1, or shFOXA1#2; group differences are assessed by one-way ANOVA. **E**, DIA protein expression analysis of SNARE related genes in shNC and shFOXA1 A549 cells. Group differences are assessed by t-test.

