## Extended data Fig. 1 for "FOXA1 preserves cell polarity and restrains lysosome biogenesis in non-small cell lung adenocarcinoma"

**A**

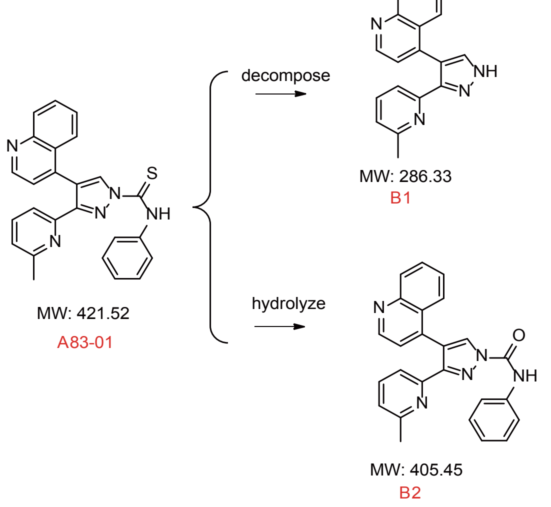


**B**

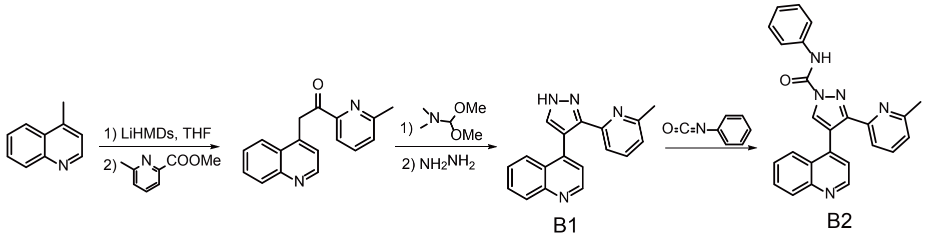


**C**

| Compound | IC50 (M) |
|---|---|
| A83-01 | 4.91e-09 |
| Dasatinib | 9.71e-08 |
| B1 | 5.54e-09 |
| B2 | 2.17e-08 |

**D**

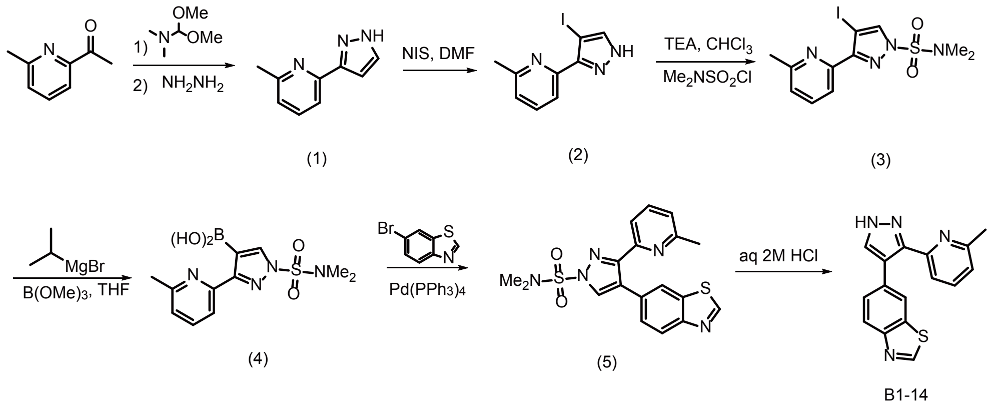


**E**

| NUMBER | NAME | ACCESSION NUMBER | ACTIVITY |
|---|---|---|---|
| 1 | TGFBR1(h) | GenBank NM_004612 | -3 |
| 2 | Abl(h) | GenBank U07563 | -1 |
| 3 | ALK4(h) | GenBank NM_004302 | -1 |
| 4 | TNIK(h) | GenBank NM_015028.1 | -1 |
| 5 | Abl (M351T)(h) | GenBank U07563 | 0 |
| 6 | Flt1(h) | GenBank AF063657 | 1 |
| 7 | Abl (H396P) (h) | GenBank U07563 | 3 |
| 8 | cKit(D816H)(h) | GenBank X06182 | 3 |
| 9 | MAP4K4(h) | GenBank NM_004834.4 | 3 |
| 10 | Flt4(h) | GenBank NM_182925 | 4 |
