## Supplementary figures and images for "FOXA1 preserves cell polarity and restrains lysosome biogenesis in non-small cell lung adenocarcinoma"

### Extended data Fig. 2

Extended data Fig. 2

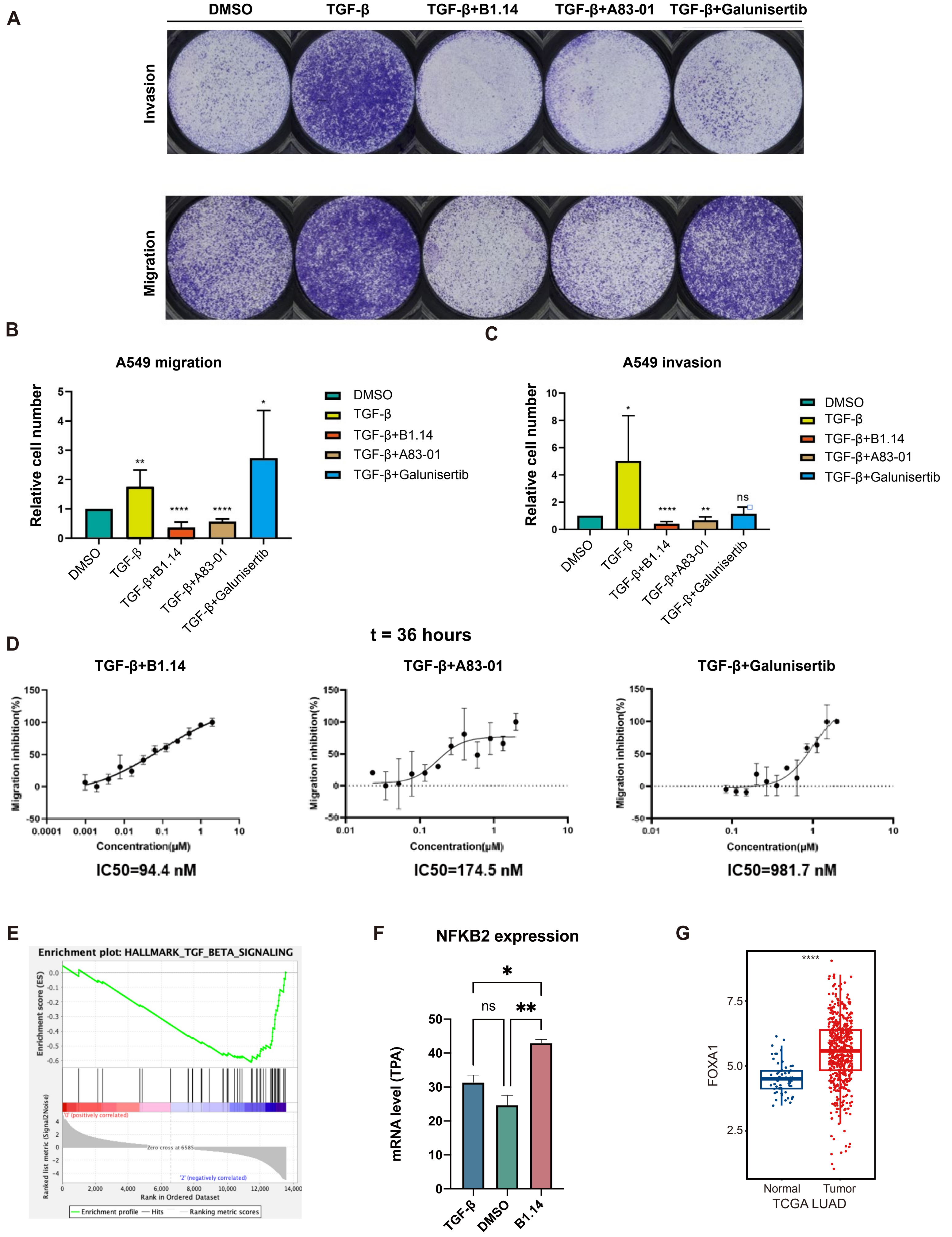

### Extended data Fig. 3

Extended data Fig. 3

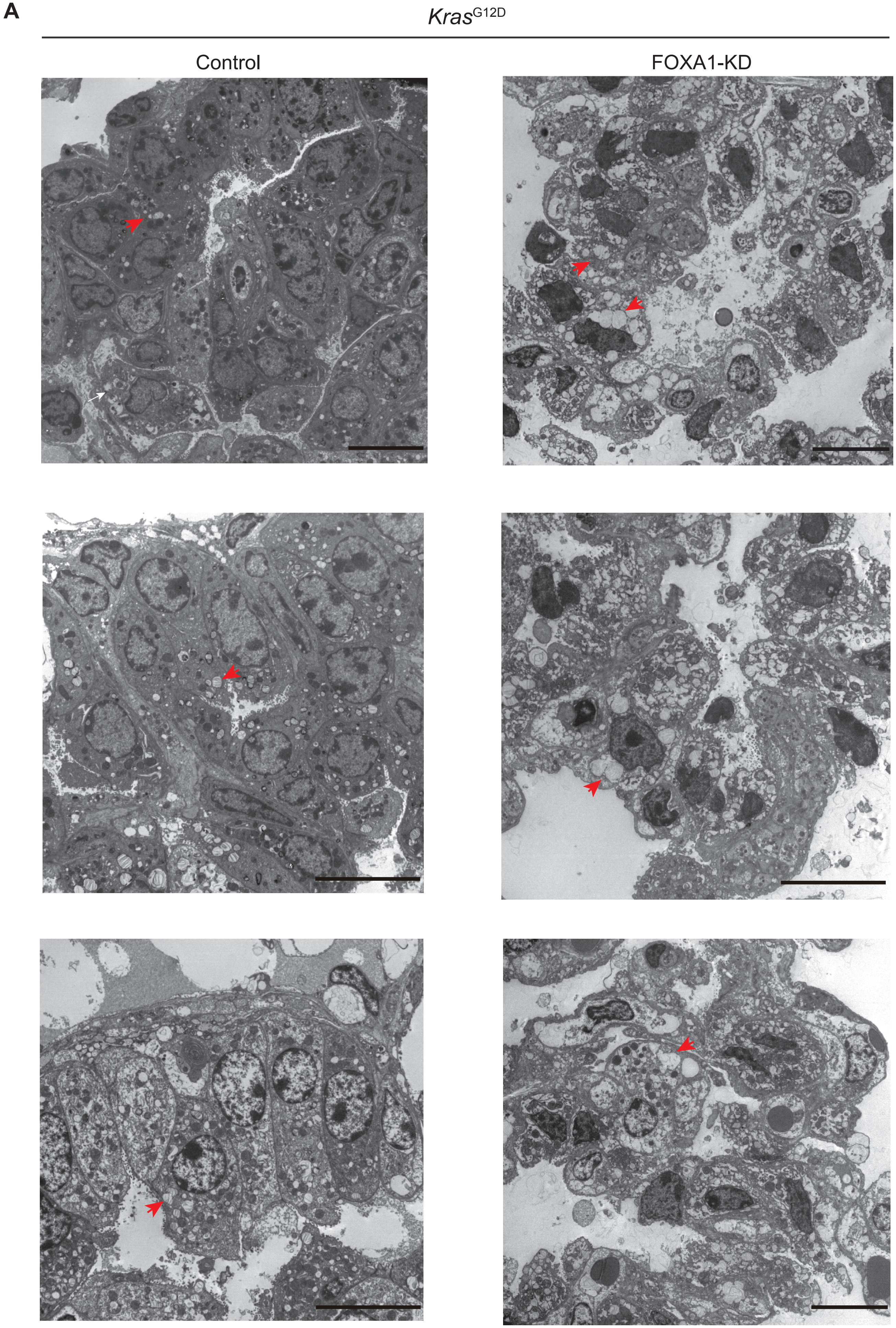


scale bar = 10 μm

### Extended data Fig. 4

Extended data Fig. 4

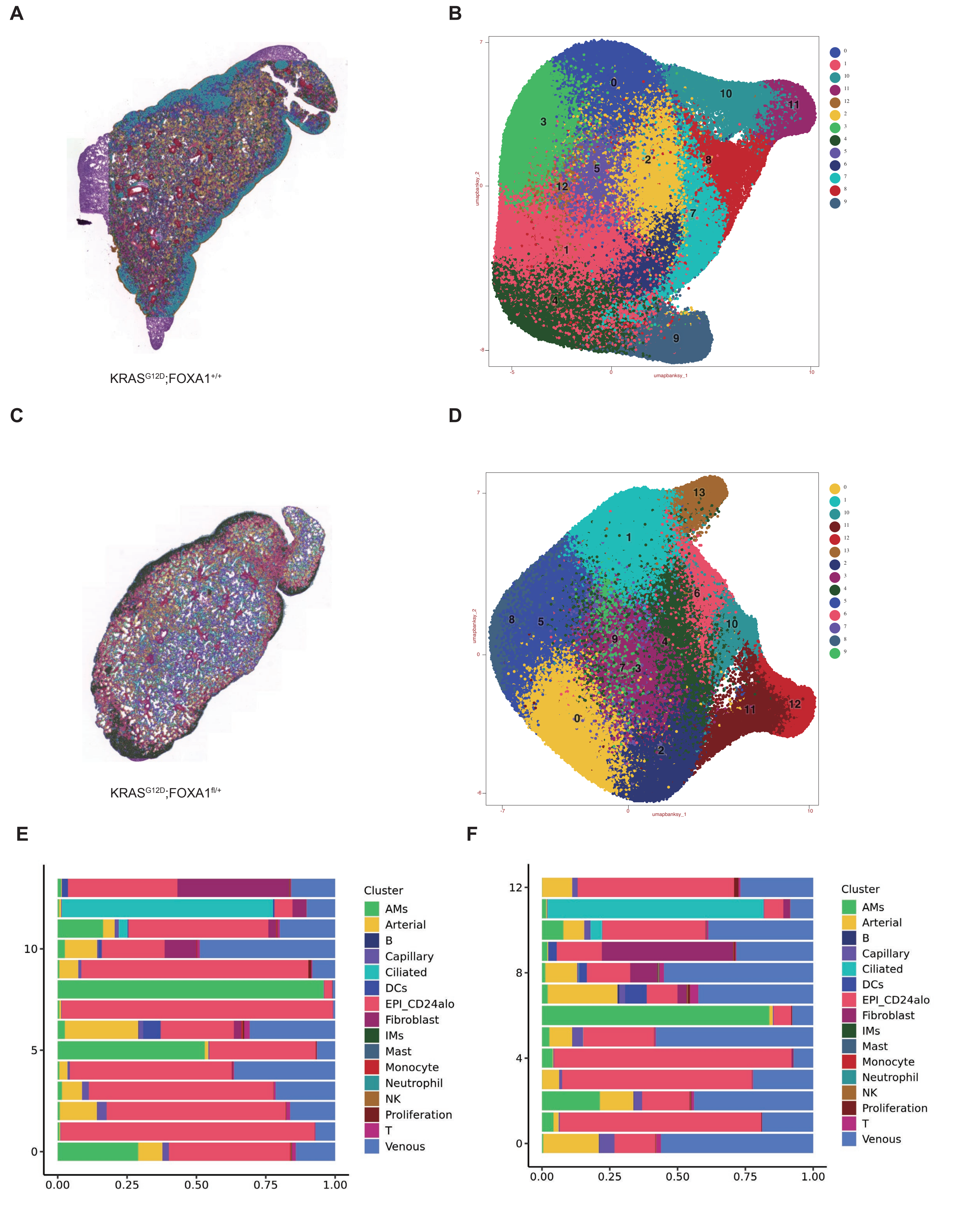

### Extended data Fig. 5

Extended data Fig. 5

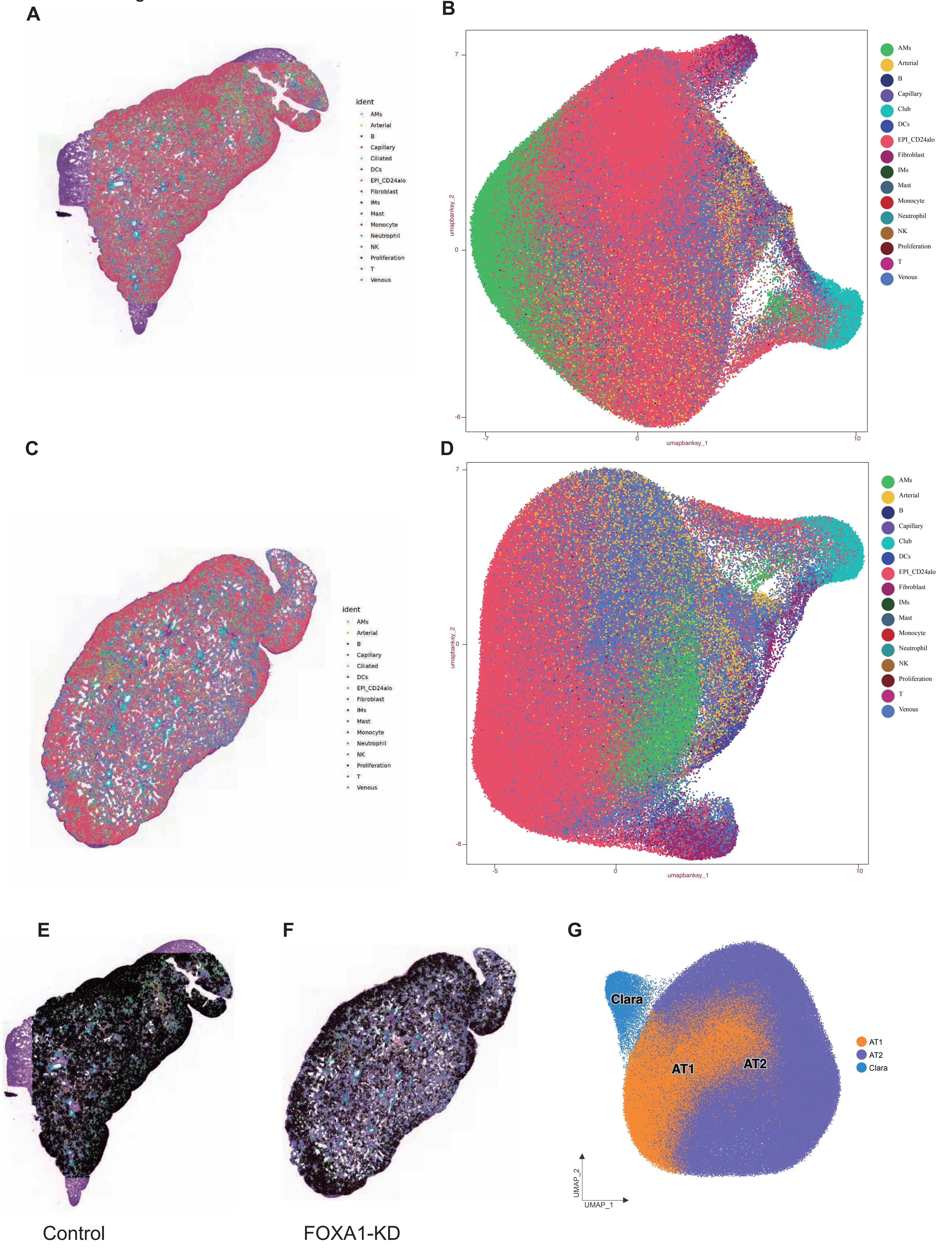

### Extended data Fig. 6

Extended data Fig. 6

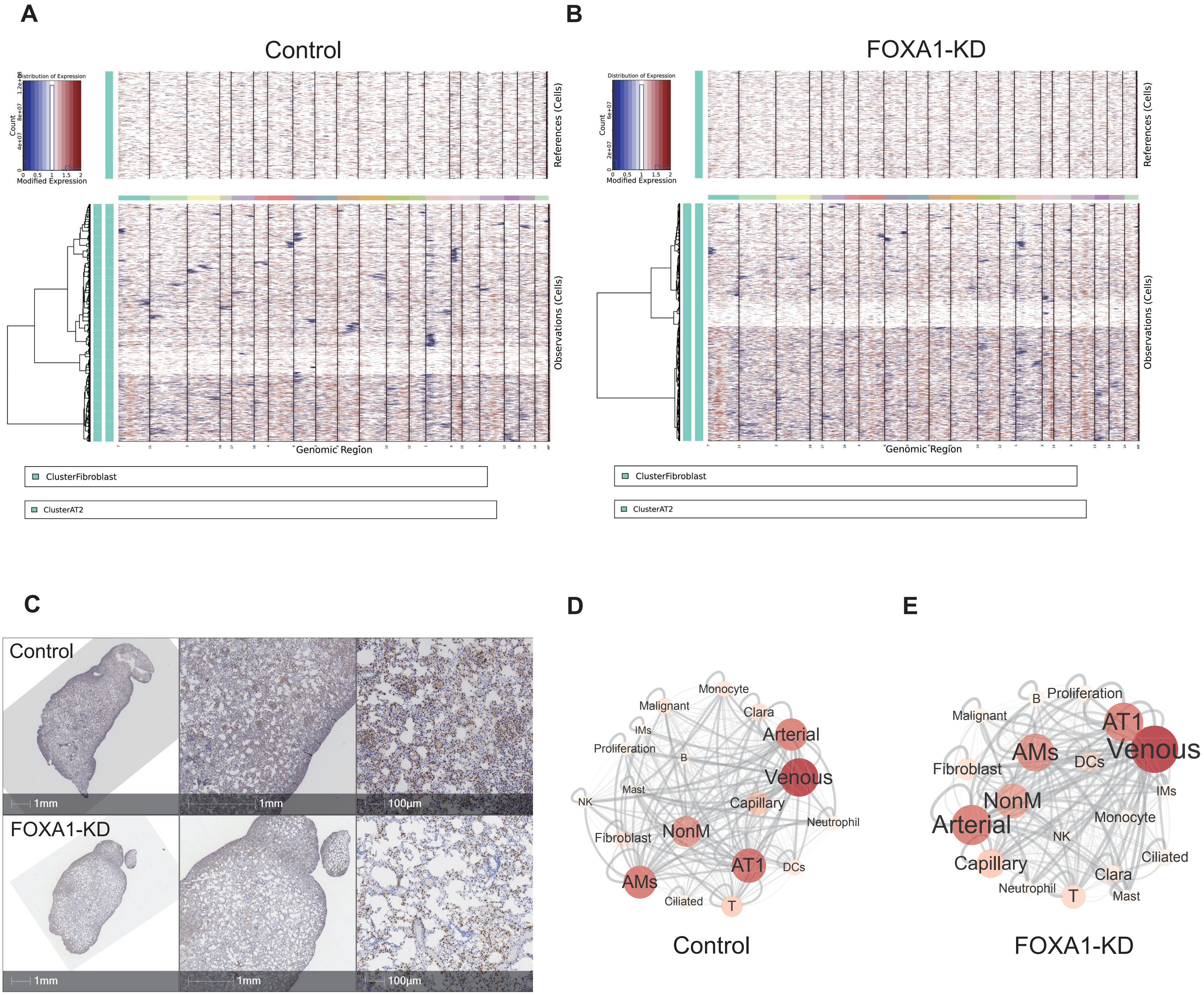

### Extended data Fig. 7

Extended data Fig. 7

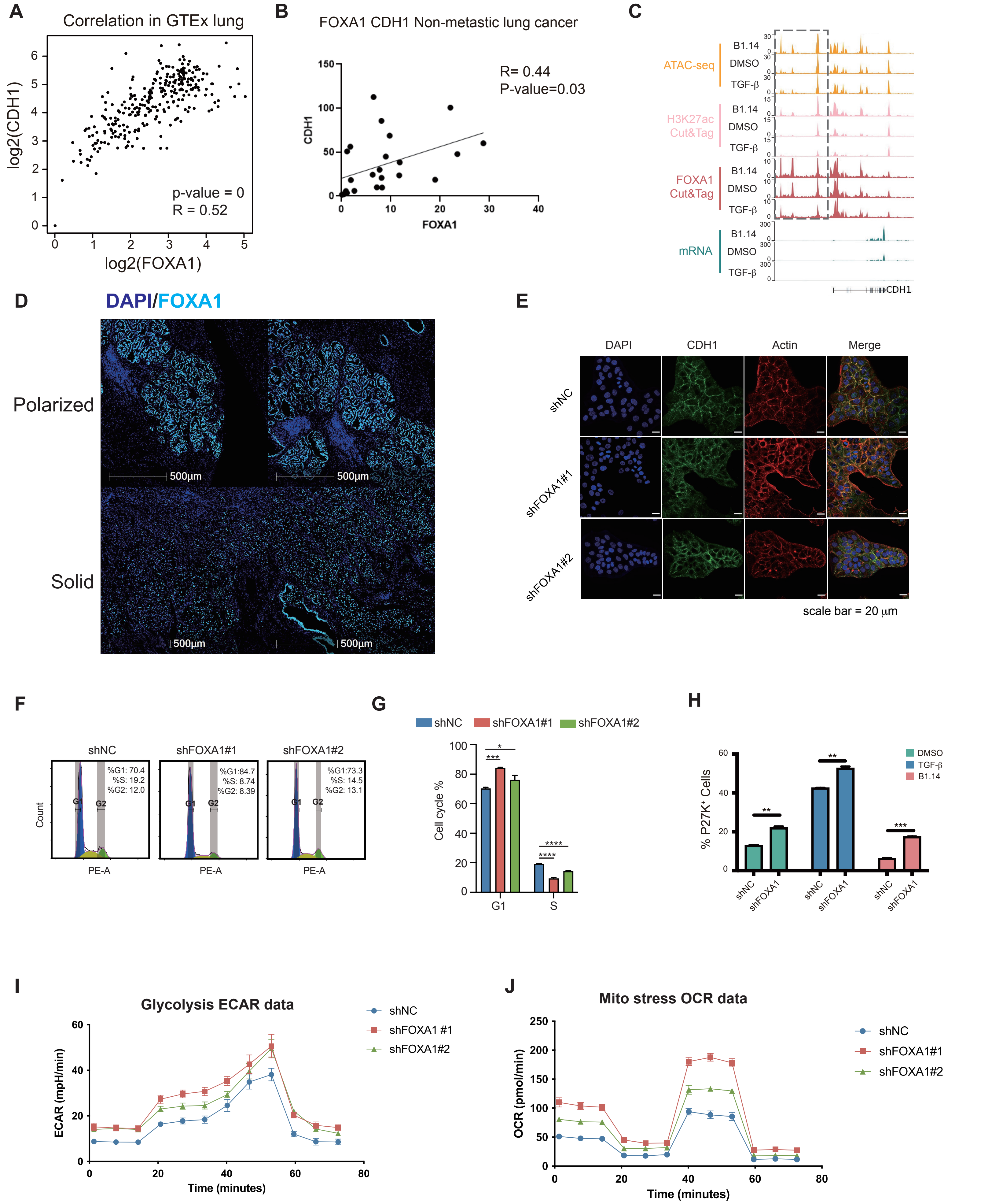

### Extended data Fig. 8

Extended data Fig. 8

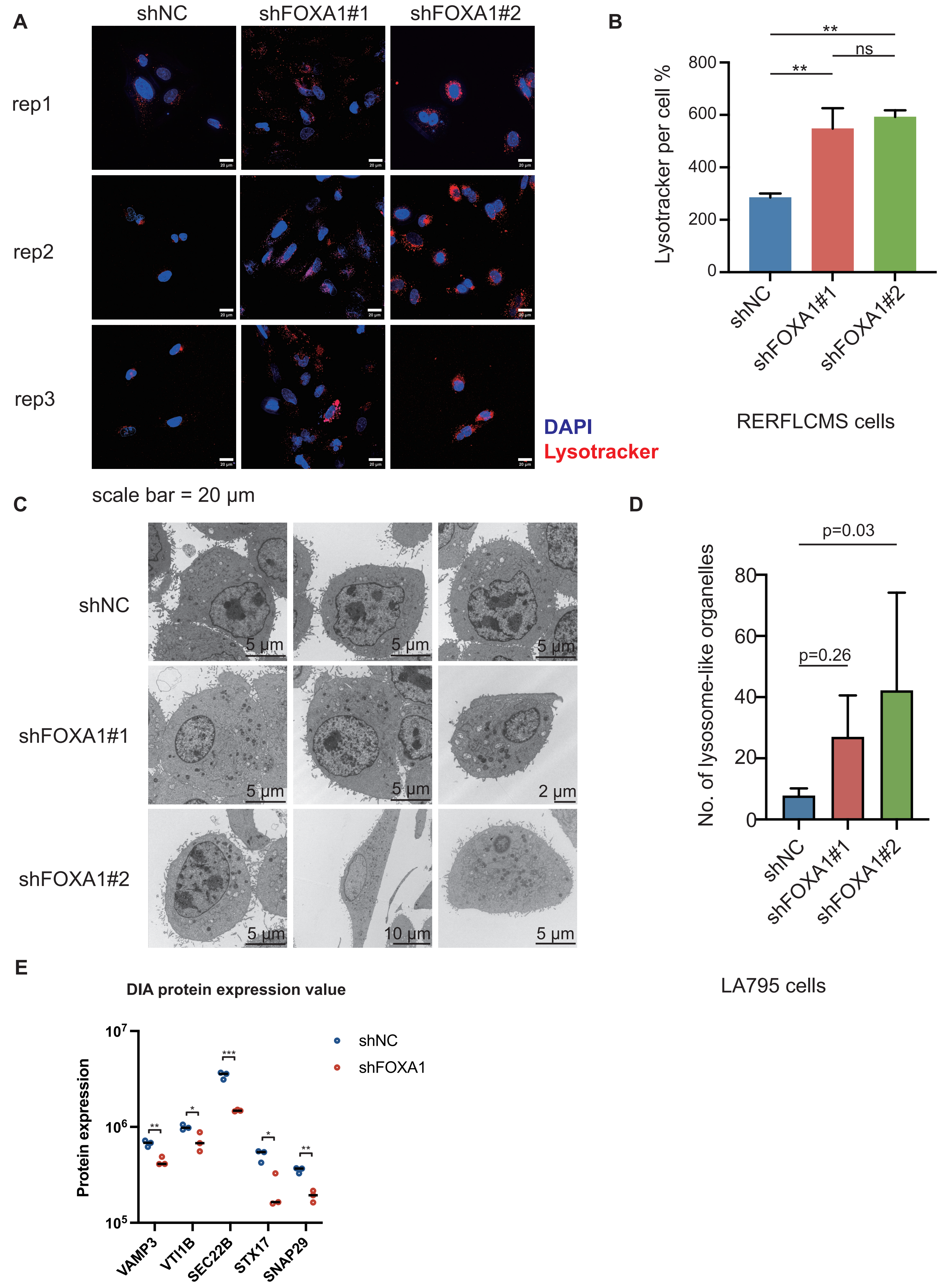
